# High-accuracy meets high-throughput for microbiome profiling with near full-length 16S rRNA amplicon sequencing on the Nanopore platform

**DOI:** 10.1101/2023.06.19.544637

**Authors:** Xuan Lin, Katherine Waring, John Tyson, Ryan M. Ziels

**Affiliations:** Civil Engineering, The University of British Columbia, Vancouver, BC, Canada; Environmental Microbiology, British Columbia Center for Disease Control Public Health Laboratory, Vancouver, BC, Canada; Pathology and Laboratory Medicine, The University of British Columbia, Vancouver, BC, Canada

## Abstract

Amplicon sequencing of small subunit (SSU) rRNA genes is a foundational method for studying microbial communities within various environmental, human, and engineered ecosystems. Currently, short-read platforms are commonly employed for high-throughput applications of SSU rRNA amplicon sequencing, but at the cost of poor taxonomic classification. The low-cost Oxford Nanopore Technologies (ONT) platform is capable of sequencing full-length SSU rRNA genes, but the lower raw-read accuracies of previous ONT sequencing chemistries have limited accurate taxonomic classification and de novo generation of operational taxonomic units (OTUs) and amplicon sequence variants (ASVs). Here, we examine the potential for Nanopore sequencing with newer (R10.4+) chemistry to provide high-throughput and high-accuracy full-length 16S rRNA gene amplicon sequencing. We present a sequencing workflow utilizing unique molecular identifiers (UMIs) for error-correction of SSU rRNA (e.g. 16S rRNA) gene amplicons, termed ssUMI. Using two synthetic microbial community standards, the ssUMI workflow generated consensus sequences with 99.99% mean accuracy using a minimum UMI subread coverage threshold of 3x, and was capable of generating error-free ASVs and 97% OTUs with no false-positives. Non-corrected Nanopore reads generated error-free 97% OTUs but with reduced detection sensitivity, and also generated false-positive ASVs. We showcase the cost-competitive and high-throughput scalability of the ssUMI workflow by sequencing 90 time-series samples from seven different wastewater matrices, generating ASVs that were tightly clustered based on sample matrix type. This work demonstrates that highly accurate full-length 16S rRNA gene amplicon sequencing on Nanopore is possible, paving the way to more accessible microbiome science.

## 1. Introduction

The amplification and sequencing of small subunit (SSU) ribosomal RNA (rRNA) genes (e.g. 16S and 18S rRNAs) is a widely used method to study the diversity and taxonomic composition of microbial communities within a variety of environments. The foundational work of Fox and Woese (1) utilized the conserved function of rRNA across all self-replicating cells to establish the first phylogenetic description of the domains of life, and provided a basis for taxonomically classifying microorganisms based on their evolutionary divergence. Since then, comparative analysis of SSU rRNA gene sequences has enabled the discovery of new uncultivated microbial lineages (2, 3), surveys of microbial community composition in host-associated (4, 5) and natural environments (6–8), and the design of oligonucleotide hybridization probes for environmental monitoring of select taxa (9, 10). Within the past decade, the throughput of SSU rRNA sequence generation has been enhanced by so-called ‘next-generation’ sequencing platforms, such as Roche 454 (8, 11) and Illumina (12) platforms, which are capable of sequencing millions of amplicons generated over hundreds of samples (13, 14).

While next-generation sequencing platforms have provided an affordable and high-throughput approach for generating SSU rRNA gene sequences from multiplexed environmental samples, the taxonomic resolution of amplicon sequences generated from such technologies is limited by their short read lengths (e.g. up to ∼500 bp in paired-end mode (15)). To circumvent this limitation, high-throughput amplicon sequencing of environmental SSU rRNA genes typically relies on the use of conserved primers to amplify discrete variable regions (e.g. V1-V9 regions in 16S rRNA). The selection of the variable region for amplification of 16S rRNA gene fragments can introduce biases into diversity estimates, as the 16S rRNA gene does not evolve evenly along its length (16). The accuracy of taxonomic assignment of 16S rRNA sequences has also been shown to increase with amplicon sequence length, with full-length 16S rRNA sequences required to capture most taxonomic ranks (17, 18). Moreover, the taxonomic characterization of SSU rRNA gene fragments from unknown microorganisms relies on their comparison to reference databases of full-length sequences. The widespread application of short-read sequencing platforms for SSU rRNA gene profiling has resulted in a decreased rate of full-length sequence generation, which is needed for phylogenetic analysis of novel lineages as well as for the development of new oligonucleotide hybridization probes (19–21). Thus, there is a need for new high-throughput approaches capable of sequencing full-length SSU rRNA gene fragments to improve taxonomic classification of microbiomes, as well as increase the number of full-length SSU rRNA sequences in public databases.

Recently, there have been advancements in single-molecule sequencing technologies capable of generating long-reads, such as the Pacific Biosciences (PacBio) and Oxford Nanopore Technologies (ONT) platforms (15). While these long-read sequencing platforms can alleviate many of the abovementioned problems associated with classifying short reads, their raw sequence data has been limited by high error rates (0.5-2%) compared to second-generation sequencers (<1%) (22–24). These high error rates of raw long-read platforms can obfuscate SSU amplicon sequence clustering and taxonomic assignments (25, 26), impacting the accuracy of such methods for microbiome profiling. As ONT sequencers (e.g. MinION) have a relatively low capital cost (27) and can be used in field settings (28–31), developing high-throughput and accurate SSU amplicon sequencing with the ONT platform could help to advance microbiome science in diverse applications worldwide.

To circumvent higher error rates in raw long-reads, previous strategies have utilized various forms of consensus sequencing approaches for error correction, which involve redundant sequencing of multiple copies of the sample DNA template molecule of interest to obtain a consensus sequence with reduced error (22, 32–34). In particular, it was recently shown that highly accurate (>99.99%) consensus sequences could be generated on the ONT platform using unique molecular identifiers (UMIs) to tag individual DNA molecules prior to amplification and sequencing (22). In this UMI-based sequencing approach, independent reads sharing the same molecular barcodes are grouped together to enable consensus sequence generation and error correction. However, a relatively high UMI subread coverage (15-25x) was necessary to reach accuracies above 99.99% (22), thus requiring a high per-sample read depth and limiting the throughput of this method for routine microbiome science involving many samples. Since the development of this UMI-based amplicon sequencing approach, new ONT sequencing chemistries and pores have been developed (e.g. ≥R10.4) with higher raw-read accuracies (35) that could increase sample throughput by improving UMI detection as well as requiring a lower sub-read coverage for a desired consensus sequence accuracy. It could also be possible that the higher raw-read accuracies of newer ONT chemistries are sufficient for high-throughput SSU sequencing alone, without the need for read error-correction.

Here, we explore the application of ONT sequencing to high-throughput and high-accuracy full-length SSU amplicon sequencing for microbiome profiling. We present a UMI-based full-length SSU amplicon sequencing workflow, termed *ssUMI*, that employs accurate quantification of starting template molecules (near full-length 16S rRNA genes) for library preparation and leverages the higher raw-read accuracy of newer ONT chemistries for stringent UMI detection and binning. We validate the ssUMI approach using two synthetic microbial community standards, and show that it improves amplicon sequence variant (ASV) and species detection compared to quality-filtered (i.e. non-error-corrected) Nanopore reads. We also demonstrate its high-throughput scalability by sequencing 90 environmental microbiome samples at a competitive per-sample cost, thus facilitating the use of ONT long-read sequencing in large-scale microbiome studies.

## 2. Results

### 2.1 Evaluating ONT raw-read accuracy with a mock microbial community standard

To determine whether raw reads generated with the ONT ‘Q20+’ chemistry (i.e. R10.4+ flow-cell) were sufficient for full-length 16S rRNA gene amplicon sequencing analysis, we first assessed the error rate distribution for 4.4M amplicon reads sequenced from the 8-species ZymoBIOMICS Microbial Community DNA Standard (Figure 1). Without any quality filtering, the raw reads had a mean accuracy of 96.5% (Figure 1B), which is insufficient to resolve species or operational taxonomic units (OTUs) at a 97% cluster identity. We found a correspondence between the expected error (EE) rate predicted from the per-base quality (Q) scores and the empirical EE rate of the raw reads (Figure 1A). Therefore, we implemented an EE-filter threshold of 1%, which filtered 93% of the raw reads and improved the mean read accuracy to 98.8% (Figure 1B).

**Figure 1.**
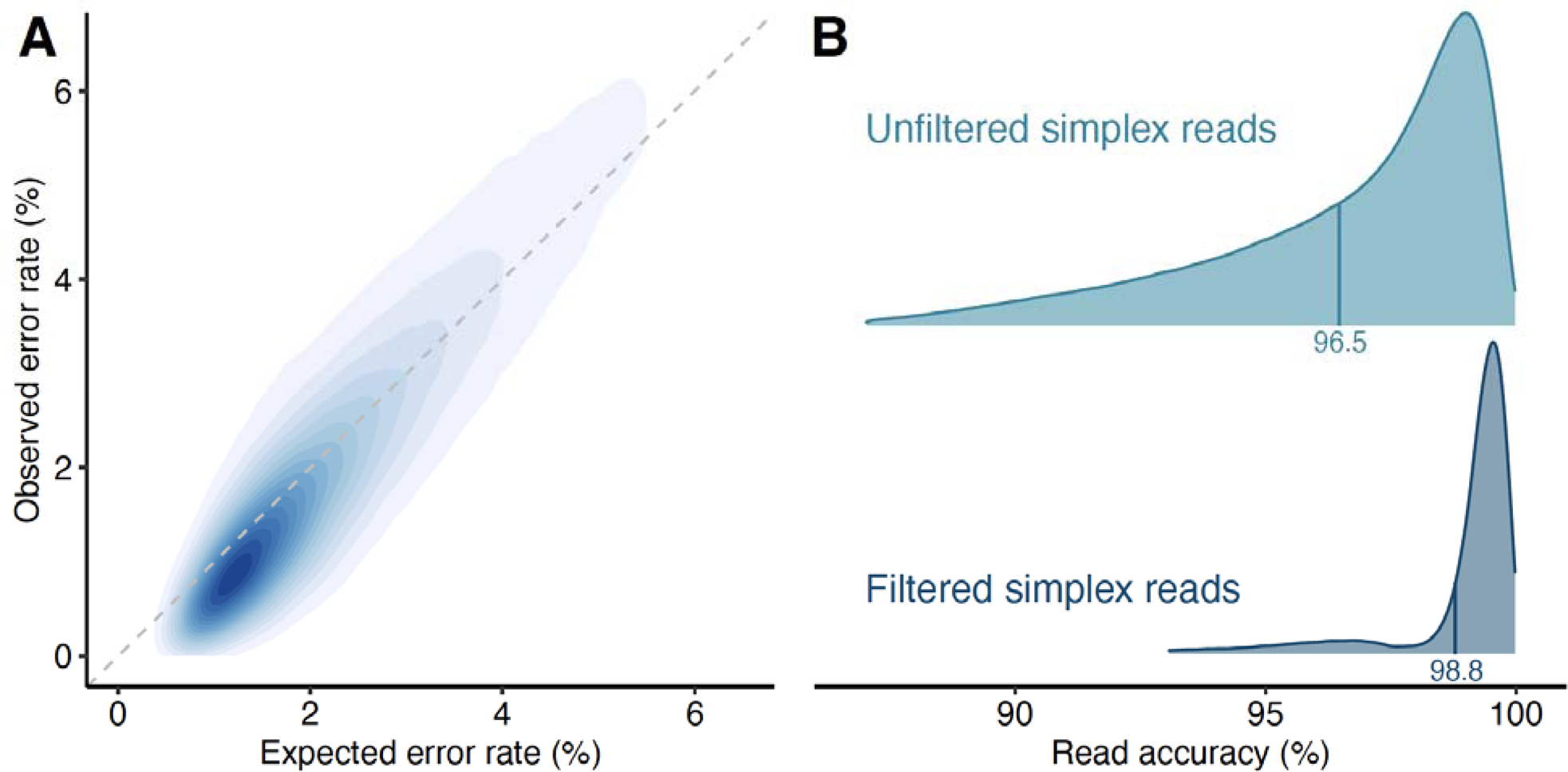
(A) Observed versus expected error (EE) rates of length-filtered raw Nanopore reads. The darker blue color indicates higher density of reads within that plot region. The dashed gray line represents a 1:1 slope. (B) Density plot of read accuracy distribution of unfiltered and length+EE-filtered raw Nanopore reads. Mean accuracy values are indicated with vertical lines, and are provided as text below the lines.

### 2.2 UMI-based amplicon sequencing of near full-length 16S rRNA genes enables accurate profiling of mock microbial communities

The above analysis of raw reads motivated us to develop a high-throughput long-read sequencing workflow for highly-accurate near full-length 16S rRNA genes on the ONT platform. We built upon the dual-UMI-based amplicon sequencing method described by Karst et al. (22), with several key modifications made here to the library preparation and data analysis steps to enable high-throughput 16S rRNA gene sequencing (Figure 2). Specifically, rather than relying on a trial-and-error approach to determine the number of starting molecules for UMI-tagging, we developed a droplet digital PCR (ddPCR) approach to accurately measure near full-length 16S rRNA gene copies within each sample. Using this approach, we determined an optimum input template molecule number of 1×10^5^ copies into the UMI-tagging PCR (Supporting Text 1), which is 10-times that used by Karst et al. (22) and is intended to increase detection of rare community members when applied to complex communities. We also developed two data analysis pipelines, termed `ssUMI_rapid` (i.e. ‘rapid mode’) and `ssUMI_std` (i.e. ‘standard mode’), that perform length filtering, EE-filtering, primer trimming, UMI detection, and consensus sequence generation using different modes of consensus read polishing (Figure 2). Due to the higher raw-read accuracy of ONT R10.4 chemistry used here, it was possible to implement more stringent UMI-based read binning in our analysis workflow by reducing the allowed UMI hamming distance to prevent erroneous read-binning. Finally, to improve the throughput of the UMI approach to 16S rRNA gene amplicon sequencing, we allowed a minimum sub-read coverage of 3x per UMI bin, rather than 15x-25x used by Karst et al. (22), to reduce the overall sequencing depth required per sample.

**Figure 2.**
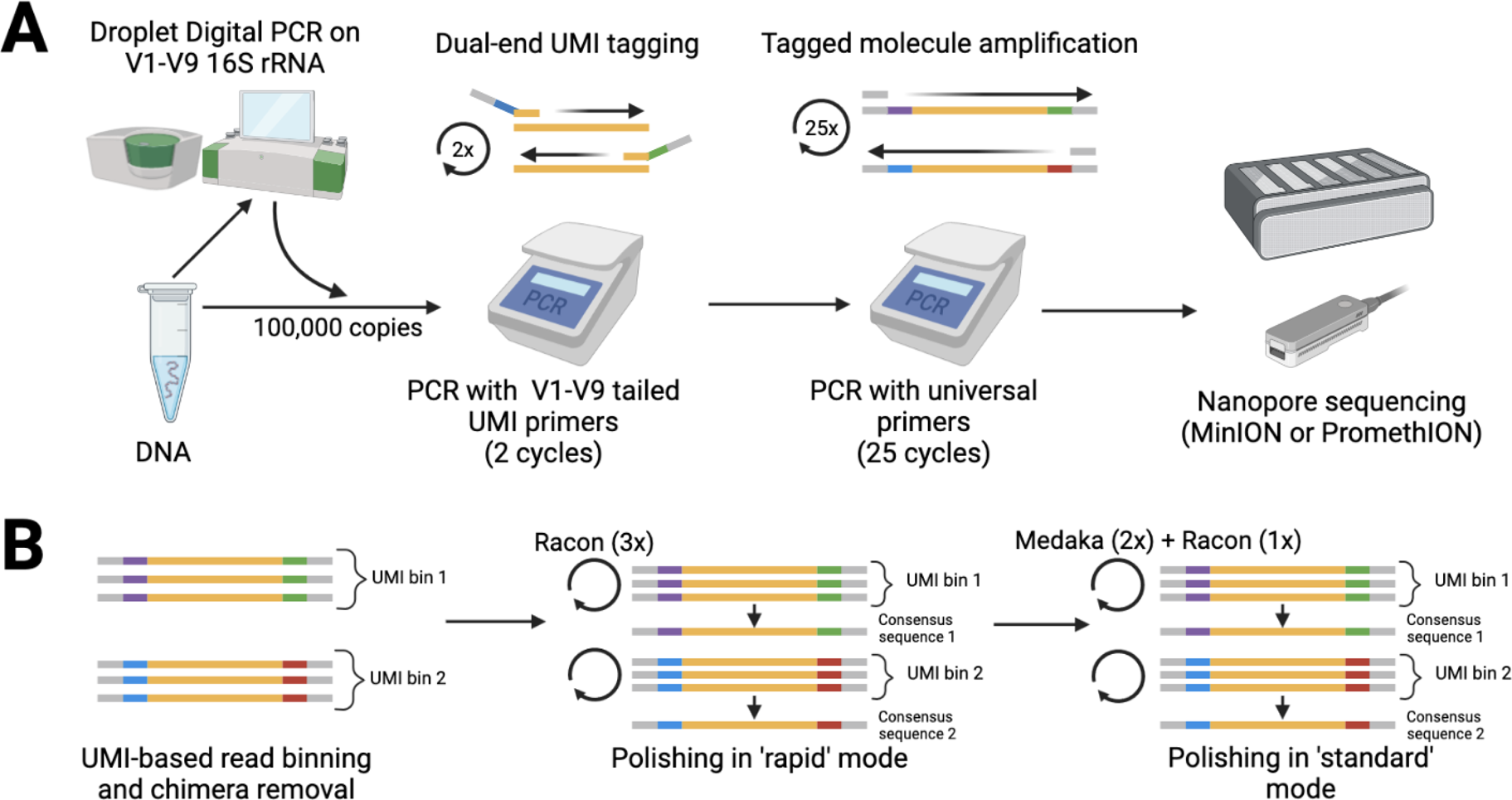
Summary of UMI-based small subunit ribosomal RNA gene sequencing (ssUMI) workflow. (A) The wet-laboratory steps, in which DNA templates are first quantified with droplet digital PCR (ddPCR) with a near full-length 16S rRNA gene assay. Then DNA containing 100,000 16S rRNA gene copies are added to the first ssUMI PCR for UMI tagging. Following two cycles of PCR, the UMI-tagged amplicons are further amplified in a second PCR using universal primers that flank the template and UMIs. After PCR amplification, the products are sample-barcoded, pooled, and sequenced on a Nanopore instrument. (B) Data analysis workflow following sequencing, in which the reads are analyzed with the ssUMI pipeline for generation of high-accuracy full-length 16S rRNA consensus sequences. Initially, reads are quality-filtered and binned based on UMIs from both ends (e.g. UMI-pairs), and chimeras are removed. Consensus sequences are polished with only Racon (3x) in ‘rapid’ mode of the workflow, or followed by Medaka (2x) and Racon again (1x) in the ‘standard’ workflow mode.

For the library generated with the 8-species ZymoBIOMICS Microbial Community DNA Standard, 8.1×10^4^ UMI-based consensus sequences (with coverage ≥3x) were generated from a single MinION R10.4 flowcell. Using only three rounds of Racon polishing, termed ‘rapid’ mode in the ssUMI workflow, the mean accuracy of the UMI consensus sequences was 99.96% (Figure 3A) and 68.1% of sequences were error-free. We found a slight increase in consensus sequence accuracy using two rounds of Medaka after the initial Racon polishing (Figure S1). Applying a final round of Racon polishing after Medaka led to a significant reduction in error rate (Figure S1), producing a mean sequence accuracy of 99.99% (Figure 3A). Interestingly, this final round of polishing with Racon after Medaka was more effective than simply applying four sequential rounds of Racon without Medaka (Figure S2). We term the three-step polishing procedure using Racon and Medaka ‘standard’ mode in the ssUMI workflow (Figure 3A). The greater consensus accuracy achieved by the ‘standard’ mode was associated with increased computational requirements compared to ‘rapid’ mode (Table S1). Notably, the mean accuracies of the UMI-based consensus sequences in both ‘rapid’ and ‘standard’ modes were higher than that of quality-filtered PacBio HiFi sequences and Illumina short-read sequences from the same Microbial Community Standard (Figure 3A). The UMI-based consensus sequences from ‘standard’ mode had a similar mean accuracy to quality-filtered synthetic long reads generated with LoopSeq (Fig. 3A), with both strategies able to generate 92.5% and 94.6% (36) error-free sequences, respectively.

**Figure 3.**
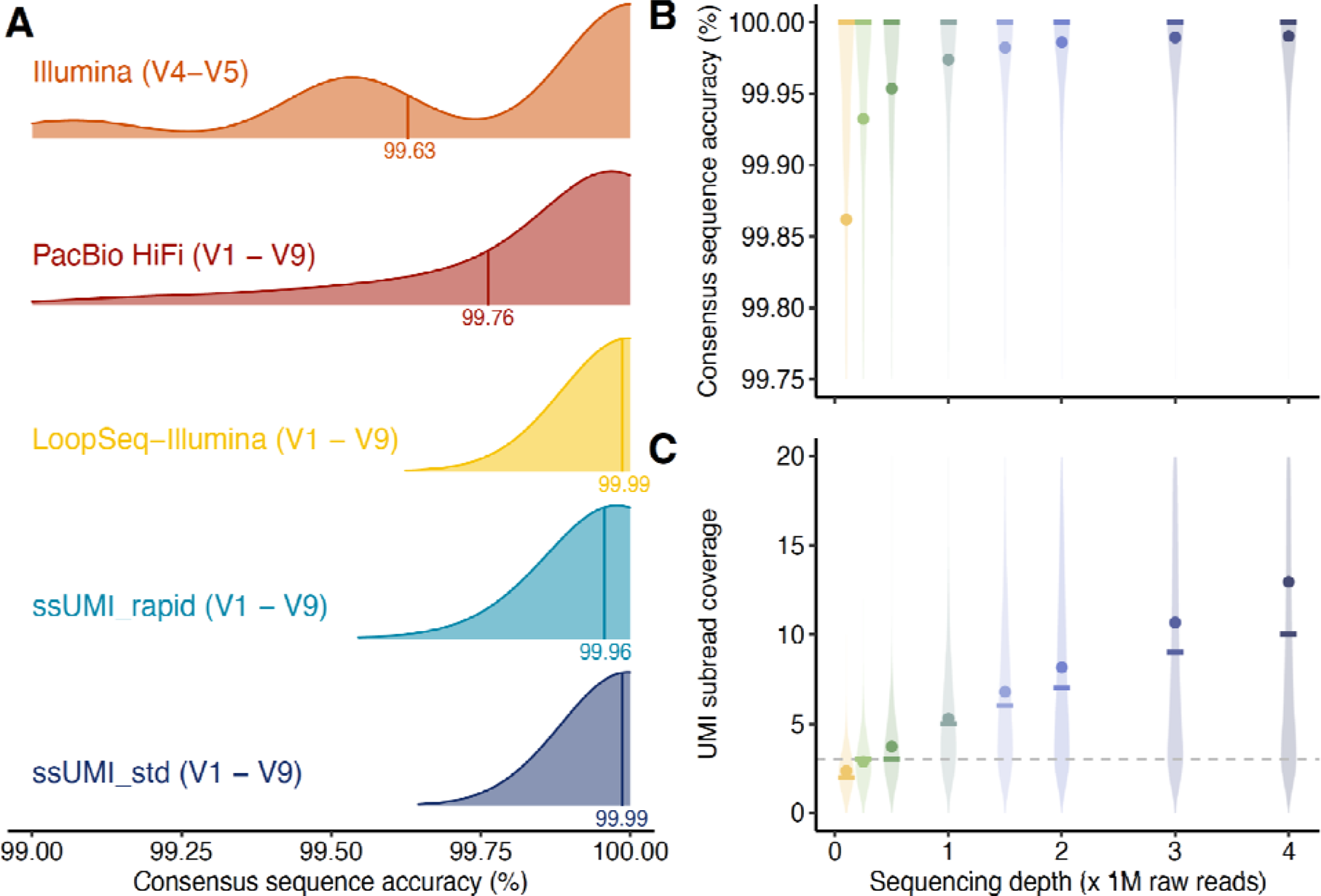
(A) Comparison of sequence accuracies obtained for the ZymoBIOMICS Microbial Community DNA Standard targeting the full-length 16S rRNA gene (V1-V9 regions) with UMI-based amplicon sequencing on ONT with `ssUMI_std` (‘standard’ mode) and `ssUMI_rapid` (‘rapid’ mode), LoopSeq (Illumina) synthetic long-reads, and PacBio HiFi sequencing, as well as short-read 16S rRNA gene amplicons (V4-V5 region) on Illumina. For all sequence data types, amplicons were quality-filtered, primer-trimmed, and contaminant sequences were removed (see Methods). Impact of raw-read sequencing depth on distribution of (B) consensus sequence accuracy distribution and (C) UMI subread coverage, for `ssUMI_std` applied to full-length 16S rRNA gene amplicon (V1-V9 regions) from the ZymoBIOMICS Microbial Community DNA Standard. Subplots B and C share an x-axis. Different colors represent raw read sequencing depths, circular points represent mean values and the crossbars represent the median values, while the shaded violin region represents the density distribution of the values at each depth. For subplot (C), the horizontal dashed line represents the minimum UMI subread coverage of 3x implemented in this study.

Because the sequencing depth of the ZymoBIOMICS Microbial Community library was much greater (e.g. 1 sample per MinION flow-cell) than would be typical in a high-throughput application with many samples, we explored the effect of per-sample raw-read depth on UMI-consensus sequence generation and accuracy. By randomly subsampling the original sequence library and generating UMI-based consensus sequences, we found a saturation-like behavior in the number of generated UMI-consensus sequences as a function of sample raw-read depth (Figure S3). As more UMI-based consensus sequences are recovered with greater raw-read depth (up to the saturation-level), this suggests that users can modify per-sample throughput based on their application and desired sensitivity for detecting rare members. We also observed that greater per-sample sequencing depth increased the UMI-based consensus sequence accuracy, which reached a mean of 99.99% above a per-sample throughput of 2M reads (Fig. 3B). This trend is largely attributed to higher consensus accuracies achieved at greater UMI-based subread coverage (22), as we also observed that UMI subread coverages increased with raw read depth (Fig. 3C). Notably, the median UMI-based consensus sequence accuracy remained at 100% and the fraction of error-free reads remained above 50% down to a per-sample raw read depth of 0.1M (Fig. 3C). At a per-sample raw read sequencing depth of 0.25M reads, the median UMI subread coverage was equal to the minimum threshold of 3x (Fig. 3C). These results indicate that per-sample raw read depth can be reduced (e.g. for sample multiplexing) while still preserving adequate UMI detection and coverage to generate highly accurate consensus sequences.

We then assessed the accuracy of ASVs and OTUs (at 97% identity) generated for the ZymoBIOMICS Microbial Community DNA Standard using both quality-filtered Nanopore raw reads and UMI-based consensus sequences. Both ‘rapid’ and ‘standard’ modes of UMI-based consensus sequences generated 100% accurate ASVs and OTUs that perfectly matched all expected 27 ASVs and 8 OTUs in the reference community (Figure 4). Surprisingly to us, quality-filtered Nanopore raw reads were also capable of generating all ASVs and OTUs perfectly from the mock community, without any false-positives (Figure 4).

**Figure 4.**
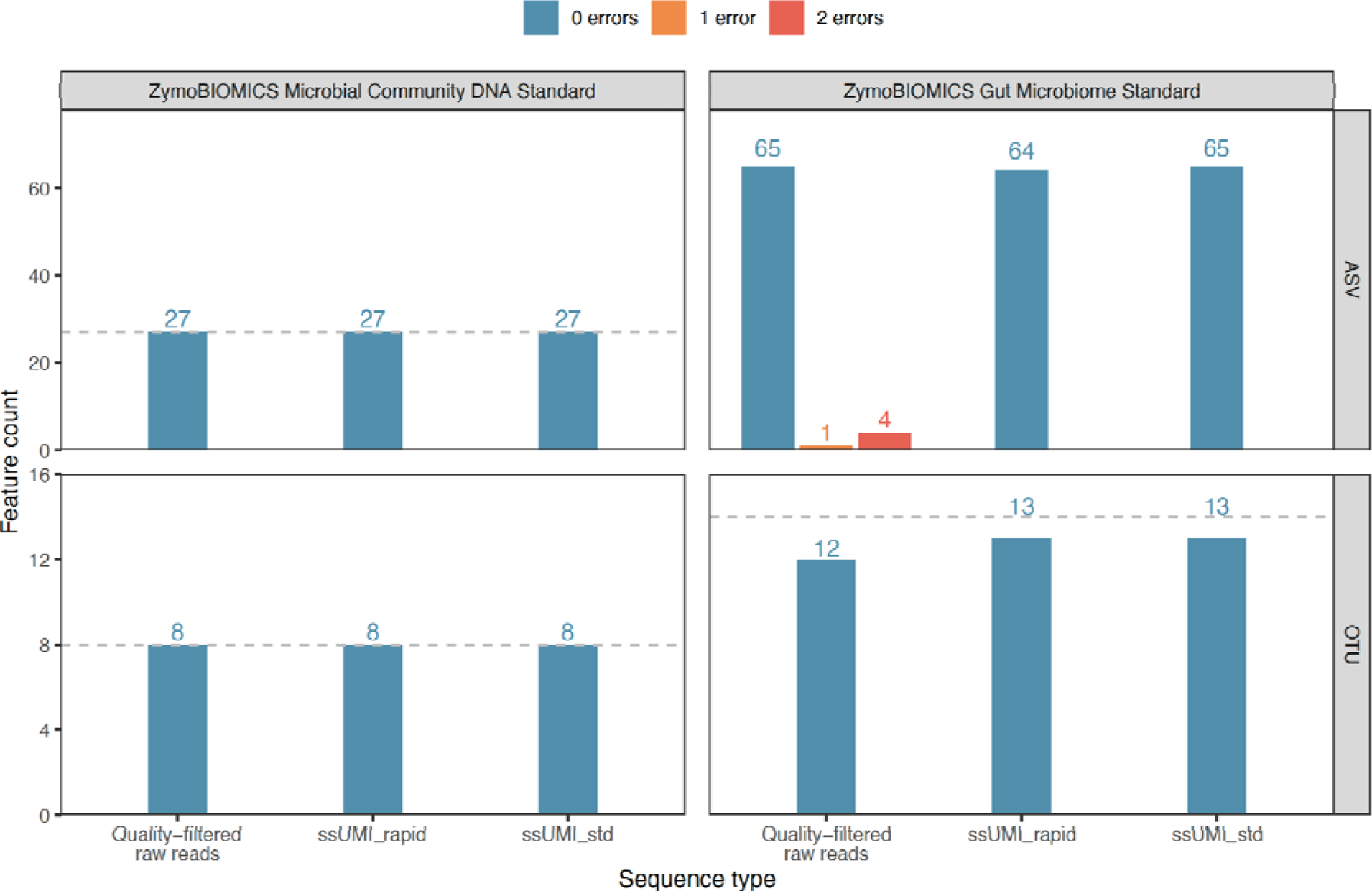
Accuracy of *de novo* near full-length 16S rRNA (V1-V9) sequence features generated for two mock microbial community standards, the 8 bacterial species ZymoBIOMICS Microbial Community DNA Standard and the 14 bacterial species ZymoBIOMICS Gut Microbiome Standard, using either quality-filtered Nanopore raw reads or Nanopore reads processed with the ssUMI pipeline in ‘rapid’ (e.g. `ssUMI_rapid`) and ‘standard’ (e.g. `ssUMI_std`) modes. The generated sequence features include amplicon sequence variants (ASVs) and operational taxonomic units (OTUs) at a 97% identity threshold. The results of the ZymoBIOMICS Microbial Community DNA Standard were generated with a single sample on a single R10.4 MinION flowcell, and that of the ZymoBIOMICS Gut Microbiome Standard were generated with 6 technical replicates of two different DNA extractions (see Methods) on two R10.4 MinION flowcells. The number of sequence errors in features are indicated with the fill colors. The dashed gray line indicates the expected number of bacterial full-length 16S rRNA features in the reference community. The lack of a dashed gray line for ASVs in the ZymoBIOMICS Gut Microbiome Standard is due to uncertainty on the true number (see Methods).

To better assess the reproducibility and capability of our ssUMI approach on a complex microbiome sample, we sequenced two DNA extracts in triplicate from the log-distributed ZymoBIOMICS Gut Microbiome Standard cell mixture containing 14 bacterial species. One DNA extract was obtained using the Qiagen MagAttract PowerSoil Pro kit and the other with a phenol:chloroform based extraction. Each extraction set was sequenced in triplicate on one R10.4 MinION flowcell, yielding 8.4 ± 1.7 × 10^5^ raw reads and 3.8 ± 0.64 × 10^4^ UMI-based consensus sequences per sequencing replicate. The measured relative abundance values in the UMI-based consensus sequences were consistent among technical replicates, and generally matched well with the theoretical abundance for community members that were more abundant than 0.01% (Figure 5A). An exception was *Lactobacillus fermentum* and *Bifidobacterium adolescentis* in the phenol:chloroform extraction, which showed a clear abundance skew that was consistent within all replicates (Figure 5A). We attribute this aberration to DNA extraction bias, rather than an artifact of the ssUMI pipeline, as this abundance skew was not observed in the replicates extracted with MagAttract PowerSoil Pro (Figure 5A). Reads from rare community members that were less than 0.01% abundance were identified sporadically with UMI-based consensus sequences. *Salmonella enterica* (0.009% theoretical abundance) was detected in all replicates of the phenol:chloroform extraction, but was only detected in the single replicate of the MagAttract PowerSoil Pro extract that had the highest number of raw reads (Figure 5A). Similarly, *Enterococcus faecalis* (0.0009% theoretical abundance) was only identified in a single technical replicate from both DNA extracts (Figure 5A).

**Figure 5.**
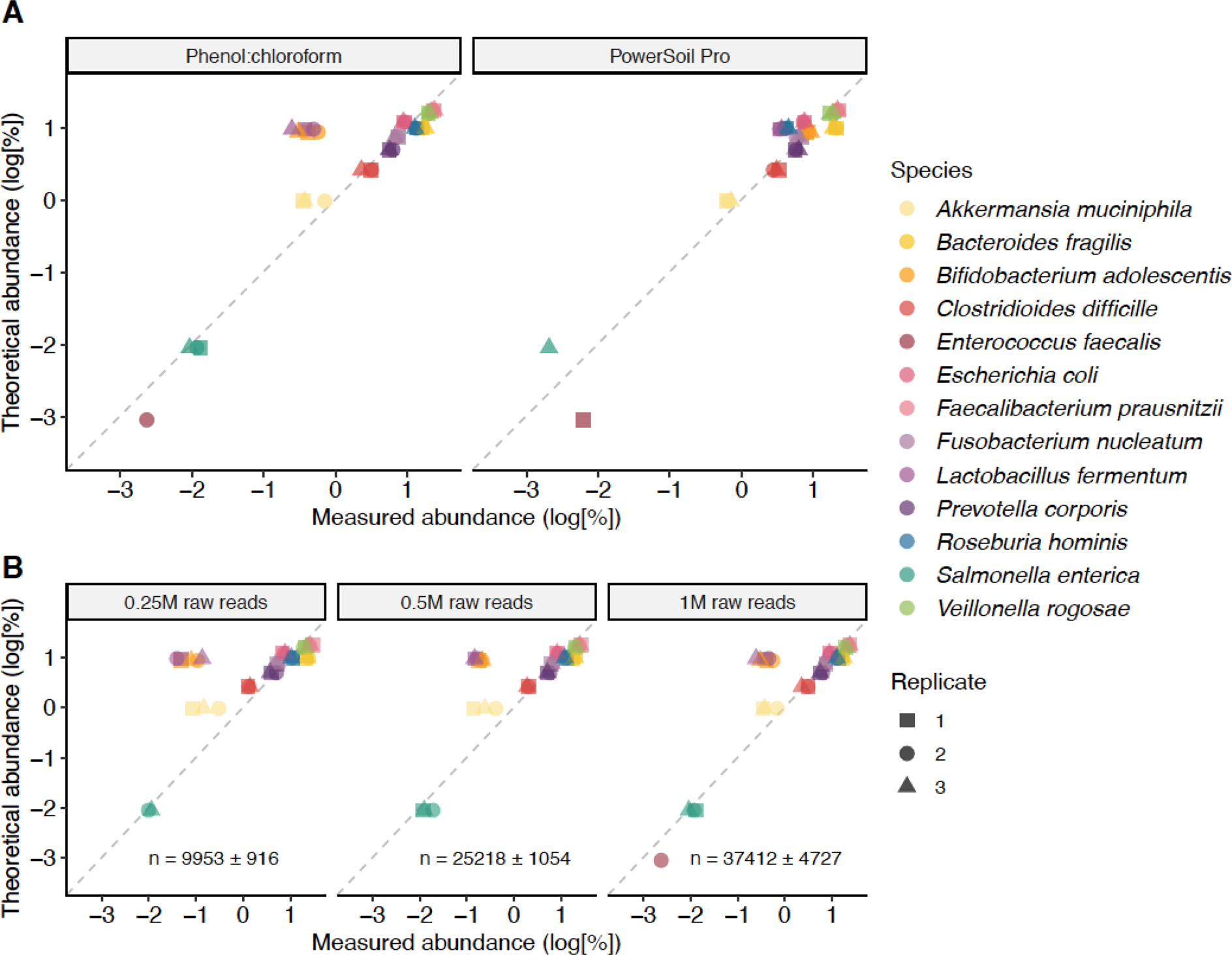
(A) Composition of ZymoBIOMICS Gut Microbiome Standard based on near full-length 16S rRNA (V1-V9 region) consensus sequences processed with the `ssUMI_std` pipeline (‘standard’ mode), in comparison to the theoretical abundances provided by the vendor. Technical PCR and sequencing replicates are shown for two different DNA extractions of the same cell mixture, phenol:chloroform and MagAttract PowerSoil Pro. (B) The impact of sample raw-read depth on the resulting microbial community composition of the ZymoBIOMICS Gut Microbiome Standard (phenol:chloroform DNA extraction) obtained with 16S rRNA consensus sequences processed with the `ssUMI_std` pipeline (‘standard’ mode). To perform this analysis, raw reads were randomly subsampled from the original sequence libraries to given depths, and UMI-based consensus sequences were generated with the ssUMI pipeline (see Methods). The text values shown in the sub-plots represent the numbers of UMI-based consensus sequences generated (+/− standard deviation of triplicates).

As increasing the sample-throughput of the ssUMI approach (i.e. more samples multiplexed in a single run) requires a reduction in the per-sample raw-read count, we investigated the effect of sample read depth on the relative abundance distribution obtained from UMI-based consensus sequences by randomly subsampling the original sequence libraries from the ZymoBIOMICS Gut Microbiome Standard phenol:chloroform DNA extraction. Reducing the raw-read depth from 1M to 0.25M did not greatly impact the observed distribution of taxa in the UMI-based consensus sequences, with the exception that detection of rare species was impacted at lower raw-read depths (Figure 5B).

For the more complex ZymoBIOMICS Gut Microbiome Standard, we recovered 65, 64, and 65 perfect ASVs from quality-filtered Nanopore raw reads, `ssUMI_rapid` consensus sequences, and `ssUMI_std` consensus sequences, respectively, by pooling sequences from all six extraction replicates (Figure 4). No errors were observed in any ASVs generated from UMI-based consensus sequences, regardless of the data analysis mode (e.g. ‘rapid’ or ‘standard’), while 5 erroneous ASVs containing one or more errors were generated with quality-filtered Nanopore raw reads (Figure 4; Tables S2-S4). No sequence type was able to recover ASVs corresponding to *Enterococcus faecalis* (0.0009% theoretical abundance) (Tables S2-S4). Only `ssUMI_std` was able to recover an ASV from *Salmonella enterica* (0.009% theoretical abundance), while this organism was missed with `ssUMI_rapid` and quality-filtered raw Nanopore reads (Tables S2-S4). After clustering into 97% OTUs, we recovered 12, 13, and 13 error-free OTUs from quality-filtered Nanopore raw reads, `ssUMI_rapid` consensus sequences, and `ssUMI_std` consensus sequences, respectively (Figure 4; Tables S5-S7). For OTUs generated with both `ssUMI_rapid` and `ssUMI_std` sequences, all bacterial species in the ZymoBIOMICS Gut Microbiome Standard were detected except for *Clostridium perfringens* (0.0002% theoretical abundance) (Tables S5 and S6). For OTUs generated with quality-filtered Nanopore raw reads, all 12 bacterial species at or above 0.01% were detected (Table S7).

### 2.3 Application of UMI based 16S rRNA amplicon sequencing for high-throughput microbiome profiling

We further demonstrated the scalability of the ssUMI workflow by applying it to 90 wastewater samples collected bi-weekly from a nearby wastewater treatment facility. A total of seven wastewater sample matrices were collected and prepared with the ssUMI workflow, and the products were sequenced on the ONT PromethION platform, generating a total of 103.8 Gb raw read data (Table S8). On average, each sample yielded 4.3 ± 1.6 × 10^5^ raw reads, with exception for three samples that were not sufficiently barcoded (Figure S4). The number of UMI-based consensus sequences increased with sequencing depth (Pearson’s r = 0.73) for all wastewater sample types (Figure S5), which is in accordance with the above-mentioned observations obtained by *in silico* subsampling of mock community libraries. Wastewater samples generated significantly more UMI-based consensus sequences than the mock community at the same sequencing depth (Student’s t-Test, *p*<0.05), indicating that the UMI-tagging and PCR was not inhibited by the complex wastewater matrices. No significant difference (ANOVA, *p*=0.135) was observed for the number of UMI-based consensus sequences generated in ‘standard’ mode from different sample types, with an average of 3.6 ± 1.1 × 10^4^ consensus sequences per sample (Fig. 6A).

**Figure 6:**
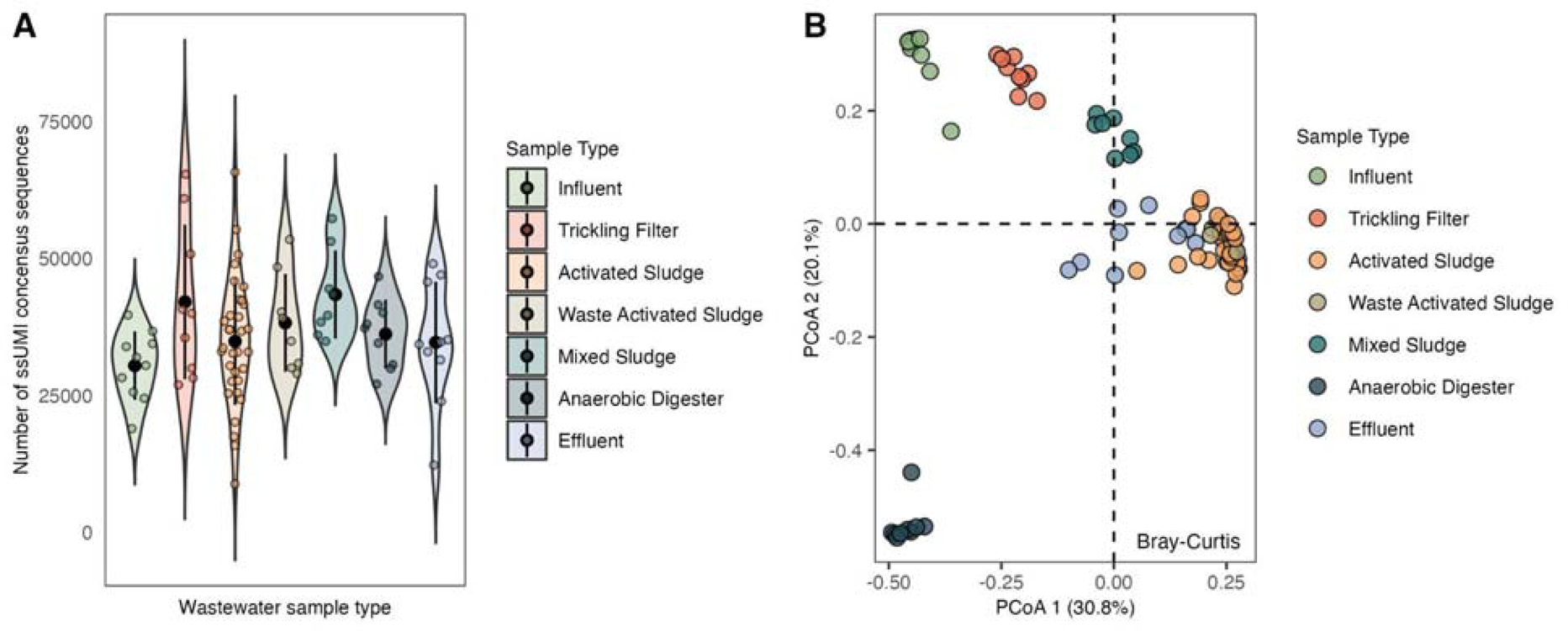
(A) Number of UMI-based consensus sequences generated with `ssUMI_std` mode for samples representing wastewater matrix types. Each colored point represents one sequenced sample, the black dots represent the mean number of consensus sequences and the bars show standard deviation. The shaded region represents the density distribution of all samples within that matrix type. (B) Principal coordinate analysis (PCoA) of ASV Bray-Curtis dissimilarity, showing the clustering of the different wastewater sample types collected over two months. Each colored point within the PCoA represents one sequenced sample.

We then generated full-length 16S rRNA gene ASVs with the UMI-based consensus sequences from all wastewater samples, yielding a total of 8349 bacterial ASVs from all sequenced samples. Based on a principal coordinate analysis (PCoA) of Bray-Curtis dissimilarity, there was a clear impact of wastewater sample type on the bacterial community structure (Figure 6B). The lowest richness (mean 1165 ± 271) was observed in anaerobic digester samples, while the highest richness (mean 3260 ± 524) was observed in mixed-sludge samples (Figure S6). Based on ASV abundance profiles, the bacterial community structures were relatively stable for each sample type over the study period (Fig. 6B; Figure S7). Influent, trickling filter, mixed sludge, and anaerobic digester samples were each tightly clustered in the PCoA, while activated sludge, waste activated sludge and effluent samples formed a relatively loose cluster (Fig. 6B).

## 3. Discussion

Numerous studies have shown the benefits of full-length 16S rRNA amplicon sequencing for taxonomic classification compared to short-read amplicon sequencing (18, 19, 37–40). However, previous approaches for high-throughput full-length 16S rRNA gene amplicon sequencing with ONT typically relied solely on alignment to reference databases (29, 41), rather than *de novo* generation of sequence features (e.g. ASVs or OTUs). NanoCLUST (42) uses k-mer based clustering of 16S rRNA gene amplicons to *de novo* generate species-level consensus sequences, but the error-profile of the consensus sequences has not been characterized and ASVs cannot be resolved with that approach. Accurate full-length 16S rRNA gene sequence features are critical for developing ecosystem specific databases (20, 43), designing organism-specific primers or probes (10, 44), and for cross-study analyses (45). In this study, we explored the potential for Nanopore sequencing to provide high-throughput and high-accuracy microbial community profiles and *de novo* generated sequence features with near full-length 16S rRNA gene amplicon sequencing. We utilized two mock microbial community standards to assess sequencing accuracy and sensitivity of non-error-corrected Nanopore reads (R10.4) alongside UMI-based consensus sequences using the ssUMI workflow.

Surprisingly, even though the mean accuracy of non-error-corrected quality filtered Nanopore raw reads was below 99%, it was still possible to generate perfect ASVs and 97% OTUs for the simpler ZymoBIOMICS Microbial Community DNA Standard. When applied to the more complex ZymoBIOMICS Gut Microbiome Standard, the non-error-corrected Nanopore reads generated false-positive (i.e. erroneous) ASVs, and the generated 97% OTUs had the poorest sensitivity of species detection among the read-types tested. The poorer sensitivity of sequence feature detection using raw Nanopore reads is likely caused by the requirement of denoising and clustering algorithms to observe a unique sequence multiple times (46), which was limited by the higher average sequence error rate. Sensitivity was also likely impacted by the large (93%) fraction of raw reads falling below the quality filter. Based on these findings, it may be appropriate to use quality filtered Nanopore raw reads for microbial profiling only if the community is known to be relatively simple with few rare members (e.g. all members >0.1% relative abundance). Future improvements in Nanopore read accuracy, such as via improved base-calling models that convert raw signal into predicted DNA sequence data (47) or by revised nanopore chemistry (48), could soon enable Nanopore reads to be used without read error-correction in diverse applications of 16S rRNA gene amplicon sequencing of complex microbiomes.

The incorporation of UMI-based error correction of 16S rRNA gene long-read amplicons via the ssUMI workflow provided higher sequence accuracy than was observed for PacBio HiFi long-reads and Illumina short-reads, and reached a similar accuracy to synthetic long-reads sequenced on Illumina. The UMI-based amplicon sequencing approach also improved the sensitivity of sequence feature detection over non-error-corrected Nanopore raw reads. The ssUMI workflow in ‘standard’ mode (`ssUMI_std`) achieved the greatest sequence accuracy, and consequently ASVs generated with these sequences had the highest sensitivity for species detection. That is, ASVs produced from `ssUMI_std` sequences included *Salmonella enterica* that was present at less than 0.01% theoretical abundance, whereas ASVs from this species were missed with `ssUMI_rapid` sequences due to residual errors reducing unique sequence counts. We therefore introduced both ‘standard’ and ‘rapid’ modes here depending on use-cases: ‘standard’ mode being more appropriate if computational resources are not limiting and/or high-sensitivity is required, and ‘rapid’ mode being more appropriate for analysis where detection of rare species is not a priority and/or when compute resources are limited. With either approach, the use of UMI-tags on both ends of the amplicons enables precise removal of chimeras (22), which are otherwise difficult to detect without molecular identifiers (49). The abundance distributions obtained by UMI-tagged 16S rRNA gene amplicons should also have reduced effects of PCR amplification bias, as UMI-based abundances are based on single molecule counting (50). Finally, by using the same full-length 16S rRNA primers for quantification by ddPCR and UMI-based amplicon sequencing, the ssUMI workflow can enable quantitative microbiome profiling using estimates of absolute microbial load, which has been shown to capture ecological trends not revealed by relative abundance values alone (51, 52).

We further explored the capacity of this highly accurate UMI-based sequencing method for high-throughput profiling of microbial communities in 90 complex environmental (e.g. wastewater) samples collected from seven different sampling locations over two months. The abundance profiles of full-length 16S rRNA ASVs generated with the ssUMI workflow were clustered according to sample matrices, providing insights into microbial community composition and colonization patterns within the treatment plant. For instance, the trickling filter process received flow from the treatment plant influent, and the bacterial communities in those sample types were closely clustered in the PCoA of ASV Bray-Curtis dissimilarity. Similarly, waste activated sludge samples represent activated sludge biomass collected from the recycle stream, and thus the co-clustering of communities from these sample types in the PCoA was also expected. The relatively high similarity between the microbial community in activated sludge and secondary clarifier effluent was likely the result of the carry-over of activated sludge microbes into treatment plant effluent. These results align with previous efforts using short-read sequencing that observed microbial immigration can impact spatial variation in microbial diversity and community structure across transects of full-scale wastewater treatment plants (53, 54). Overall, our results showcase that the UMI-based ONT 16S rRNA gene consensus sequences can be used for multi-sample experiments to gather ecological insights into microbial dynamics.

At a target per-sample sequencing depth of 0.5M raw reads, sufficient to generate ∼40×10^3^ UMI-based consensus sequences, we find that 16S rRNA gene amplicon sequencing with the ssUMI workflow on ONT is cost-competitive with other current sequencing platforms (Table S9). However, this cost landscape will inevitably change as long-read and short-read platforms continue to progress in terms of accuracy and throughput. Regardless, unlike other existing sequencing platforms, 16S rRNA gene amplicon sequencing on ONT platforms (e.g. PromethION P2 or MinION) can easily be performed in a typical laboratory, thus providing more flexibility and faster data turnaround. Overall, the findings of this study indicate that UMI-based 16S rRNA gene sequencing on the Nanopore platform can be applied in a high-throughput manner to sensitively and accurately measure microbial community structures in complex microbiomes. This development should help to democratize microbiome science in laboratories and field settings worldwide.

## 4. Materials and Methods

### 4.1 Sources of DNA

*Escherichia coli*, Strain 83972 (BEI Resources, Manassas, VA, USA) and two mock community products (Zymo Research, Irvine, CA, USA) were used for validation of the ssUMI pipeline. DNA from *E. coli* was extracted with the MagAttract HMW DNA kit (Qiagen, Hilden, Germany) following the manufacturer’s protocol for DNA extraction from Gram-negative bacteria. The ZymoBIOMICS Microbial Community DNA Standard (cat. no.: D6306, lot no: 213089; Zymo Research, USA) is a DNA standard consisting of eight evenly distributed bacteria (3 gram-negative and 5 gram-positive) species. The ZymoBIOMICS Gut Microbiome Standard (cat. no: D6331, lot no: ZRC194753; Zymo Research, USA) is a cell standard comprising 18 bacterial strains (14 species), 2 fungal strains, and 1 archaeal strain mixed at log-distributed cell concentrations. DNA was extracted from 125 ul of fully resuspended ZymoBIOMICS Gut Microbiome Standard cell mixture with a phenol:chloroform extraction protocol (55, 56) and the MagAttract PowerSoil Pro DNA kit (Qiagen, USA), following the published protocol or manufacturer’s instructions, except for the following modifications: 1) MetaPolyzyme treatment (57) was used for cell lysis for the phenol:chloroform extraction; and 2) the volume of MagAttract Suspension G beads and Buffer QSB1 were doubled for the MagAttract PowerSoil Pro DNA kit.

Primary clarifier effluent (referred to herein as “influent”), trickling filter effluent, activated sludge mixed liquor, waste activated sludge, mixed primary and secondary sludge (“mixed sludge”), anaerobic digester sludge, and secondary clarifier effluent (“effluent”) samples were collected from a wastewater treatment facility in the Vancouver area (British Columbia, Canada) from June to August 2022. Wastewater samples were shipped to the University of British Columbia (British Columbia, Canada) on ice, aliquoted, and concentrated via flocculation (influent, trickling filter, activated sludge, waste activated sludge, and effluent) or centrifugation (mixed sludge and anaerobic digester sludge) within 24 hr (see Supporting Text), and stored at −20°C before extraction. The preserved wastewater samples were thawed at 4°C, and DNA was extracted with the MagAttract PowerSoil Pro DNA kit (Qiagen, Hilden, Germany) using an Opentrons-2 (Opentrons Labworks, Queens, NY, USA) automated liquid handler (Supporting Text). Extracted DNA samples were quantified with Qubit™ dsDNA HS Assay Kit using a Qubit 4 fluorometer (Invitrogen, Waltham, MA, USA).

### 4.2 Molecule tagging and PCR amplification

DNA templates were first quantified with a droplet digital PCR full-length 16S rRNA assay (see Supporting Text). The UMI-tagging and PCR amplification were conducted with DNA containing 100000 16S rRNA gene copies per reaction (based on ddPCR quantification), using a modified PCR program and conditions from ONT (Custom PCR UMI protocol) (58). In brief, 16S rRNA genes were dual-tagged with UMIs in 2 cycles of PCR (ssUMI-PCR1), then amplified with two additional PCR runs (ssUMI-EarlyPCR2 and ssUMI-LatePCR2) consisting of 10 and 15 cycles, respectively. Each ssUMI-PCR1 reaction contained 5 ul diluted DNA template (100000 16S rRNA copies by ddPCR), 500 nM UMI-containing forward (8F) and reverse (1391R) primers (Integrated DNA Technologies, Coralville, IA, USA) targeting the 16S rRNA gene (see Table S10), and 25 ul 2x Platinum™ SuperFi™ II Green PCR Master Mix (Thermo Fisher Scientific, Waltham, MA, USA) in a 50 ul total volume. The two secondary rounds of PCR (i.e. ssUMI-EarlyPCR2 and ssUMI-LatePCR2) were comprised of 18 ul cleaned PCR products from the previous step, 100 nM forward and reverse universal primers, 1mM MgCl_2_, and 25 ul 2x Platinum™ SuperFi™ II Green PCR Master Mix in 50 ul total reaction volumes. The ssUMI PCR thermocycling conditions were optimized for the full-length 16S rRNA gene to reduce non-specific amplification and chimeras by keeping a low PCR cycle number and using a longer extension time (Table S11). All primer sequences and detailed thermocycling conditions for each PCR are summarized in Tables S10 and S11. After each PCR step, PCR products were cleaned with Mag-Bind® TotalPure NGS beads (0.6x beads/sample ratio, Omega Bio-tek, Norcross, GA, USA) following the manufacturer’s instructions.

### 4.3 Sequencing Library Preparation and Sequencing

Nanopore sequencing libraries of UMI-tagged full-length 16S rRNA amplicons from the ZymoBIOMICS Microbial Community DNA Standard, ZymoBIOMICS Gut Microbiome Standard, and wastewater samples were prepared using the ONT Ligation Sequencing Kit 12 and Native Barcoding Kit (SQK-LSK112, SQK-LSK112.24, and SQK-LSK112.96, respectively) following the manufacturer’s instructions, and sequenced in three different runs for 72 hours: 1) ZymoBIOMICS Microbial Community DNA Standard was sequenced on a MinION R10.4 flowcell; 2) the three technical replicates of ZymoBIOMICS Gut Microbiome Standard extracted with phenol:chloroform extraction and MagAttract PowerSoil Pro were barcoded and sequenced on two MinION R10.4 flowcells; 3) wastewater samples (90 in total) and a no-template control were barcoded and sequenced on two PromethION R10.4 flowcells.

The ZymoBIOMICS Microbial Community DNA Standard was also prepared for Illumina sequencing of V4-V5 regions of the 16S rRNA gene (non-UMI-tagged) with primers 515F-Y 5’-GTGYCAGCMGCCGCGGTAA-3’, and 926R 5’-CCGYCAATTYMTTTRAGTTT-3’ (59), following the Earth Microbiome Project 16S Illumina Amplicon Protocol (13) at the Biofactorial (Bio!) high-throughput facility (University of British Columbia, BC, Canada) using an Illumina MiSeq in 2×300 paired end mode.

### 4.4 Nanopore read basecalling and processing

Nanopore sequencing raw data was first base-called with guppy v6.3.8 using the super high accuracy model (dna_r10.4_e8.1_sup.cfg), and then demultiplexed using guppy v6.3.8 with default settings.

For assessment of Nanopore raw reads (i.e. without UMI-based error correction) for microbial profiling of the ZymoBIOMICS Microbial Community DNA Standard and ZymoBIOMICS Gut Microbiome Standard, raw reads were length-filtered (1200 - 2000 bp) and quality filtered based on a maximum expected-error (EE) rate of 1% with VSEARCH v11 (60) using the ‘`--fastq_filter`’ command (60). Nanopore sequencing adapters and 16S rRNA primers (8F/1391R) were removed from the unfiltered and quality filtered raw reads with Porechop v0.2.4 (https://github.com/rrwick/Porechop) and cutadapt v2.7 (61), as previously described (22). Sequences not containing both primers were discarded.

Quality filtered raw reads were de-replicated using USEARCH (v11.0) (64) with the ‘`-fastx_uniques`’ command and a minimum number of sequence observations of 2. Amplicon sequence variants (ASVs) were generated with the de-replicated sequences using the UNOISE3 algorithm (65) in USEARCH (v11.0), with a minimum unique size of 10. Operational taxonomic units (OTUs) clustered at 97% identities were generated from size-sorted sequences de-replicated with the ‘`-cluster_otus`’ command in USEARCH (v11.0).

### 4.5 ssUMI data processing pipeline

Raw reads from each sample were analyzed with the ssUMI data-analysis pipeline for generation of high-accuracy consensus sequences. The ssUMI pipeline was derived from the `longread_umi` package developed by Karst et al. (22), which identifies UMI sequences in raw reads, bins raw reads by shared UMI-pairs (e.g. UMIs from both ends), and generates consensus sequences for each UMI bin. We made several key modifications to adapt the workflow for the newer ONT sequence chemistry applied to 16S rRNA gene amplicon sequencing. Specifically, a new EE-rate based quality filtering was applied to the raw reads using VSEARCH v11 with the ‘`--fastq_filter`’ command and an EE threshold of 10%. Length filtering for our near full-length 16S rRNA target was modified to 1200 to 2000 bp. Due to the lower raw-read error rates for Nanopore reads generated with the R10.4 chemistry, the allowed error rate in a 36 bp UMI-pair (e.g. 18 bp UMIs from each end) was reduced from 6 bp to 4 bp, the maximum mean errors per UMI-pair in a bin was set to 2, and the minimum allowed UMI cluster size was reduced to 3. We also implemented two modes of consensus sequence generation, termed ‘`ssUMI_rapid`‘ for fast analysis and ‘`ssUMI_std`‘ for high-accuracy analysis. In `ssUMI_rapid` mode, sequences were polished with a 3 rounds of Racon v. 1.4.10 (62); while in `ssuMI_std` mode, sequences were polished with a three-step method, consisting of 3 rounds of Racon, followed with 2 rounds of Medaka v.1.7.2, and a final round of Racon. The `‘r104_e81_sup_g610’` model was used for polishing with Medaka. Minimap2 v2.17 (63) was used for mapping raw reads during polishing steps, with the ‘`-ax map-ont`’ flag. Scripts associated with the ssUMI data processing pipeline are available at: https://github.com/ZielsLab/ssUMI.

ASVs and 97% OTUs were generated with UMI-based consensus sequences following the same procedure described above for quality filtered raw Nanopore reads.

To investigate the effects of sequencing depth, we randomly subsampled raw reads from mock community samples using seqtk (66) to specified read counts. We then processed the sub-sampled reads with the ssUMI analysis workflows described above.

### 4.6 PacBio, LoopSeq, and Illumina read processing

PacBio HiFi (CCS) reads of amplicons targeting the full rRNA operon of the ZymoBIOMICS Microbial Community DNA Standard (cat. D6306, lot no: ZRC190811) (22) were downloaded from the European Nucleotide Archive under accession ERR3813246. PacBio HiFi sequences were quality-filtered using VSEARCH v11 with the ‘`--fastq_filter`’ command and an EE-rate threshold of 1%. Sequences were trimmed to the same length as the 16S rRNA amplicons generated in this study by truncating the reads at the 8F and 1391R primer sequences using cutadapt v2.7, as described above.

Synthetic long-reads of amplicons targeting the V1-V9 region of 16S rRNA genes from the ZymoBIOMICS Microbial Community DNA Standard (cat. D6306, Lot ZRC190811) generated with the LoopSeq 16S rRNA Kit and Illumina 2×150 bp sequencing (36) were downloaded from NCBI under BioProject PRJNA644197. LoopSeq amplicons were quality-filtered using VSEARCH v11 with the ‘`-- fastq_filter`’ command and an EE-rate threshold of 1%. Sequences were trimmed at the primers used for their generation (forward: “AGAGTTTGATCMTGGC”; reverse: “TACCTTGTTACGACTT”) (36) using cutadapt v2.7, as described above.

Illumina V4-V5 amplicons were processed using DADA2 pipeline (67). Only forward reads were included in the analysis. Quality filtering of forward reads were conducted with the ‘`filterAndTrim`’ function using ‘`trimLeft=15, truncLen=230, maxN=0, maxEE=1`’ arguments.

### 4.7 Characterizing sequence accuracy and abundances with microbial community standards

For the ZymoBIOMICS Microbial Community DNA Standard, curated reference rRNA operon sequences (16S-23S rRNA) were obtained from Karst et al. (22). Reference 16S rRNA gene sequences were retrieved using barrnap (https://github.com/tseemann/barrnap), trimmed to the 8F/1391R primer sequences using cutadapt v2.7, and were de-replicated using USEARCH (v11.0) with the ‘`- fastx_uniques`’ command. For the ZymoBIOMICS Gut Microbiome Standard, the genome assembly and polishing approaches were not adequately described by the vendor, and therefore we downloaded their provided reference genome assemblies for prokaryotic members [RefSeq Accessions: GCA_028743295.1, GCA_028743435.1, GCA_028743335.1, GCA_028743755.1, GCA_028743555.1, GCA_028743355.1, GCA_028743375.1, GCA_028743635.1, GCA_028743535.1, GCA_028743315.1, GCA_028743095.1, GCA_028743735.1, GCA_028743275.1, GCA_028743415.1, GCA_028743255.1, GCA_028743395.1, GCA_028743455.1, GCA_028743775.1, GCA_028743475.1], concatenated the scaffolds, and polished the assembly using PacBio HiFi reads from three metagenomes of the same mock community (NCBI Accession: PRJNA680590) using one round of Racon. Reference 16S rRNA gene sequences were retrieved and de-replicated as described above. Based on this workflow, we detected 66 unique bacterial 16S rRNA gene sequences in the ZymoBIOMICS Gut Microbiome Standard. However, `ssUMI_std` sequences generated 65 error-free ASVs without the detection of two low-abundant bacterial species (Figure 4; Table S1); therefore the true number of bacterial 16S rRNA ASVs in this community is uncertain (Figure 4).

To assess sequence accuracy of different datasets, reads were mapped to their corresponding 16S rRNA gene reference databases using minimap2 v2.17 with the ‘`-- cs`’ flag, and the mapping statistics were filtered using samtools (68) with ‘`view -F 2,308`’. The minimap2 flag ‘`-ax sr`’ was used for mapping Illumina short-reads, ‘`-ax map-ont`’ for mapping raw and UMI-corrected Nanopore reads, and ‘`-ax map-pb`’ was used for mapping PacBio HiFi reads. Mapping files were parsed in R v4.2.1 as previously described (22), and error rates were calculated as the sum of mismatches, insertions, and deletions divided by the alignment length. Reads mapping to contaminants (see below) were filtered before summarizing sequence accuracies. Read mapping files were also parsed to investigate the relative read abundances based on total sum scaling of species within the ZymoBIOMICS Gut Microbiome Standard using UMI-based consensus sequences (Figure 5).

Following previous guidelines used for characterizing sequence accuracy of mock communities with ambiguous reference genomes and closely-related strains (38), we manually confirmed the sequence accuracy of ASVs and 97% OTUs generated with the ZymoBIOMICS Gut Microbiome Standard that did not match our reference sequences by querying them with BLASTn against the NCBI nr database. If an ASV or OTU sequence matched a 16S rRNA gene with 100% identity and 100% query cover from a species that is present in the Microbiome Standard, it was considered a true positive sequence (Table S12). Otherwise, the sequence feature was assigned the error rate observed via read mapping above. If an ASV or OTU sequence matched a species from a different genus than was present in the Microbiome Standard at >97% identity and 100% query cover, this was considered a contaminant (Table S13). Reads that mapped to contaminant ASVs or OTUs were filtered when characterizing the error rate profiles of Nanopore raw reads, UMI-based consensus sequences, PacBio HiFi reads, and Illumina short-reads.

### 4.9 Wastewater sample analysis

Reads from wastewater samples were processed with the ssUMI workflow in ‘standard’ mode. To compare the ssUMI PCR efficiencies for wastewater samples and mock communities, a Student’s t-Test was performed on the number of ssUMI consensus sequences generated with wastewater samples and ZymoBIOMICS Microbial Community DNA Standard with the same sequencing depth. Wastewater samples were binned into four group based on the sequencing depths (e.g. 0.1-0.2M, 0.2-0.3M, 0.3-0.7M, 0.7-1.3M raw reads), and the average number of ssUMI consensus sequences in each group was compared to the ones generated for the mock community with 0.1M, 0.25M, 0.5M, and 1.0M raw reads, respectively. The overall performance of ssUMI workflow for different wastewater sample matrices were further examined with ANOVA on the number of Nanopore raw reads and ssUMI consensus sequences.

Microbial community analysis of wastewater near full-length 16S rRNA ASVs was conducted using the vegan package in R v4.2.1 (69). The alpha diversity was assessed with species richness and Shannon index. Beta diversity was estimated using principal coordinate analysis (PCoA) with Bray-Curtis dissimilarity.

### 4.10 Data availability

The sequencing data for this project, including both ZymoBIOMICS mock microbial community standards, are available in the NCBI under BioProject PRJNA974480. Accessions for individual wastewater samples are provided in Table S8.

## Acknowledgements

We would like to thank Metro Vancouver staff, including Parisa Chegounian, and David Blair, for assisting with wastewater sample collection. We would also like to thank Mads Albertsen for his helpful insights on this work. This work was funded by a Natural Science and Engineering Research Council of Canada (NSERC) Alliance Grant (556792–2020) to R.M.Z.

