## Supporting Information for "High-accuracy meets high-throughput for microbiome profiling with near full-length 16S rRNA amplicon sequencing on the Nanopore platform"

**This PDF file includes:**

Supporting Text

Figures S1 to S10

Tables S1 to S15

SI References

**Supporting Information Text**

**Supplemental Text 1: Droplet digital PCR (ddPCR) assay for near full-length 16S rRNA genes.**

We developed a long-amplicon ddPCR assay for accurately quantifying total full-length 16S rRNA gene copies. This assay was developed with DNA extracted from an *Escherichia coli* isolate in order to determine the optimal input molecule number for UMI-tagging (i.e. ssUMI-PCR1). The assay was then employed to estimate the number of target molecules within DNA samples, to appropriately dilute them to the target input for UMI-tagging and PCR.

***DNA extraction and quality check for Escherichia coli isolate***

DNA was extracted from 5ml *E. coli* culture preserved with 10% glycerol in TE buffer. The culture was thawed at 4 °C for 1.5 hours, then centrifuged at 5000 x g for 10 min at room temperature to pellet the cells. The supernatant was removed and DNA was extracted from the pellet using the MagAttract HMW DNA kit (Qiagen, Hilden, Germany) following the manufacturer’s protocol for Manual Purification of High-Molecular-Weight Genomic DNA from Gram-Negative Bacteria. The extracted high-molecular-weight DNA was further incubated at 56°C for 10 minutes to encourage DNA relaxation. Extracted DNA was quantified with Qubit™ dsDNA BR Assay Kit using Qubit 4 fluorometer (Invitrogen, Waltham, MA, USA).

***Development of near full-length 16S rRNA gene ddPCR assay***

The near full-length 16S rRNA gene assay was developed on a QX200 AutoDG Droplet Digital PCR System (Bio-Rad, Woodinville, WA, USA). The ddPCR reaction was comprised of 900 nM forwarded and reverse primers (Table S10), 250 nM 515F probe with FAM reporter (Integrated DNA Technologies, Coralville, IA, USA) (Table S10), and 10 uL ddPCR Supermix for Probes (No dUTP) (Bio-Rad, Woodinville, WA, USA) in a 20 uL reaction. The number of input molecules to ddPCR was limited to 10000 molecules or less per ddPCR reaction, according to the manufacturer's recommendation. Due to the nature of long amplicons targeted in this assay, the extension time was increased to 4 minutes, together with a 50-cycle program to achieve sufficient amplification per droplet in order and reach a higher fluorescence intensity that would allow for better separation between fluorescent (i.e. ‘positive’) and non-fluorescent (i.e. ‘negative’) droplets. PCR thermocycling conditions for the near full-length 16S rRNA gene ddPCR assay are given in Table S14. Clear separation of positive and negative droplets were observed in the full-length 16S rRNA gene ddPCR assay run with a series dilution of *E. coli* genomic DNA (Figure S8), and the ddPCR-measured full-length 16S rRNA gene concentrations were linearly correlated R^2^=0.999) with the theoretical concentrations (Figure S9).

We further explored the effect of increasing input molecule numbers on the accuracy of our full-length 16S rRNA gene ddPCR assay. We increased the input molecule numbers from the recommended optimal number of 10000 molecules to 20000, 50000, and 100000 16S rRNA gene molecules. Good separation of positive and negative droplets were observed for input molecules up to 50000 16S rRNA gene copies (Figure S10). Based on this observation, we thereafter used 50000 16S rRNA gene copies for ddPCR measurements of DNA samples in the ssUMI workflow.

***Determining the optimal number of input 16S rRNA genes for UMI-tagging in the ssUMI workflow***

The optimal input 16S rRNA gene copies for UMI-tagging in the ssUMI workflow was determined based on UMI-tagging rates assessed with ddPCR. Five different *E. coli* DNA dilutions with increasing 16S rRNA gene copies (i.e. 10000 copies, 50000 copies, 100000 copies, 500000 copies, and 1000000 copies per ssUMI PCR-1) were used to determine the optimal input molecule number to the UMI-tagging step. The ssUMI tagging reaction (i.e. ssUMI-PCR1) was conducted for each *E. coli* DNA dilution in triplicate, and the total 16S rRNA gene copies after the 2-cycle ssUMI PCR-1 were measured using the full-length 16S rRNA ddPCR assay described above. The UMI-tagging efficiency was estimated as:

$UMI tagging efficiency = \sqrt{\frac{ddPCR measured total 16S rRNA gene copies in ssUMI PCR1 product}{4 \times ddPCR measured total 16S rRNA gene copies in input DNA template}}$	(1)

Significantly higher UMI-tagging efficiencies were observed with 100000 and 500000 16S rRNA gene copies than all other input molecule numbers (ANOVA and Student’s t-Test, p<0.05), but no significant difference was observed between these two treatments (Table S15). We selected 100000 full-length 16S rRNA gene copies as our optimal input molecule number for the ssUMI workflow, as higher numbers of input molecules would require greater per-sample sequencing depth to recover UMI-based consensus sequences with coverage >3x.

**Supplemental Text 2: Wastewater sample storage and DNA extraction protocols.**

***Wastewater sample storage protocol***

Primary clarifier effluent (influent), trickling filter effluent, activated sludge mixed liquor, waste activated sludge, mixed primary and secondary sludge (mixed sludge), anaerobic digester, and secondary clarifier effluent (effluent) samples were collected from a wastewater treatment plant in Vancouver area (British Columbia, Canada) in sterilized 125 mL HDPE sampling bottles, and shipped on ice to the laboratory in the University of British Columbia. Influent, trickling filter, activated sludge, waste activated sludge and effluent samples were preserved with a flocculation protocol^1^. Samples were first mixed by shaking, then aliquoted into 15 mL (influent, trickling filter, and effluent), 3 mL (activated sludge), or 1.2 mL (waste activated sludge) aliquots in triplicate. Ferric nitrate (1% (v/v), 2M) was added to each aliquot, and the pH was adjusted to 5.5-6 with 5 N sodium hydroxide. Samples were shaken to mix, and incubated at 4°C overnight for the flocculation process, then centrifuged at 12,000 x g for 20 min at 4°C. The supernatants were decanted, and finally the biomass pellets were resuspended with 150 uL Zymo DNA/RNA shield (Zymo Research, Irvine, CA, USA) then stored at -20°C until DNA extraction.

Mixed sludge and anaerobic digester samples were stored using a centrifugation protocol. Samples were first mixed by shaking, then aliquoted into 0.5 mL and 0.8 mL aliquots for mixed sludge and anaerobic digester samples, respectively. The aliquoted samples were centrifuged at 12000 *x g* for 20 min at 4°C. The supernatants were then decanted, and finally the pellets were resuspended with 150 uL Zymo DNA/RNA shield then stored at -20°C until DNA extraction.

***DNA extraction of wastewater samples***

Wastewater DNA was extracted using the MagAttract PowerSoil Pro kit with an Opentrons-2 (Opentrons Labworks, Queens, NY, USA) automated liquid handler following the manufacturer’s instructions. Samples stored at -20°C were thawed at 4°C overnight, then the whole pellet mixture was transferred to a PowerBeads Tube (Qiagen, Hilden, Germany) for cell lysis, with a batch size of 16 samples per run. As per recommendation by the manufacturer, the bead-beating time was increased to 13 minutes to provide sufficient cell lysis. The supernatant of cell lysate was then transferred to 1.5 mL Eppendorf DNA LoBind tubes (Eppendorf, Hamburg, Germany) for protein precipitation with solution CD2. All centrifugation steps were performed at 15000 *x g* for 1 min at room temperature. Steps 1 to 7 of the manufacturer’s protocol were conducted in individual tubes manually, and starting from Step 8, the DNA cleanup was performed on the Opentrons-2 liquid-handler in 2 mL deep-well 96-well plates (NEST Biotechnology, Wuxi, Jiangsu, China). The Opentrons-2 was programmed to clean the cell lysate with magnetic beads, then transfer the cleaned DNA into a new 96-well PCR plate (Bio-Rad, Woodinville, WA, USA) for downstream applications.

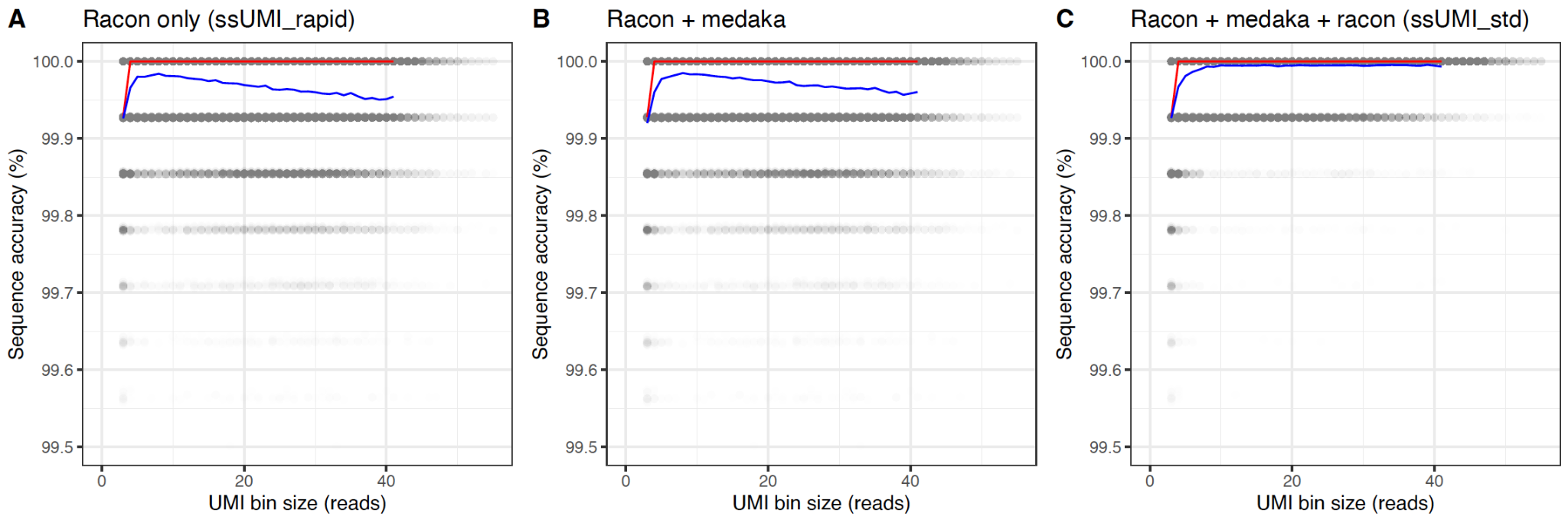

**Fig. S1.** Accuracy of UMI-based 16S rRNA gene consensus sequences (y-axis) versus UMI bin size (number of sub-reads sharing same UMI-pair; x-axis) after (A) 3-rounds of polishing with Racon (i.e. ssUMI_rapid mode); (B) 3-rounds of polishing with Racon followed by 2 rounds of polishing with Medaka; (C) 3-rounds of polishing with Racon followed by 2 rounds of polishing with Medak followed by 1 round of polishing with Racon (i.e. ssUMI_std mode). The UMI-tagged 16S rRNA gene amplicons were generated from the ZymoBIOMICS Microbial Community DNA Standard. Black points represent the accuracy of individual consensus sequences, the red line represents the median accuracy of all consensus sequences at that UMI bin size, and the blue line represents the mean sequence accuracy.

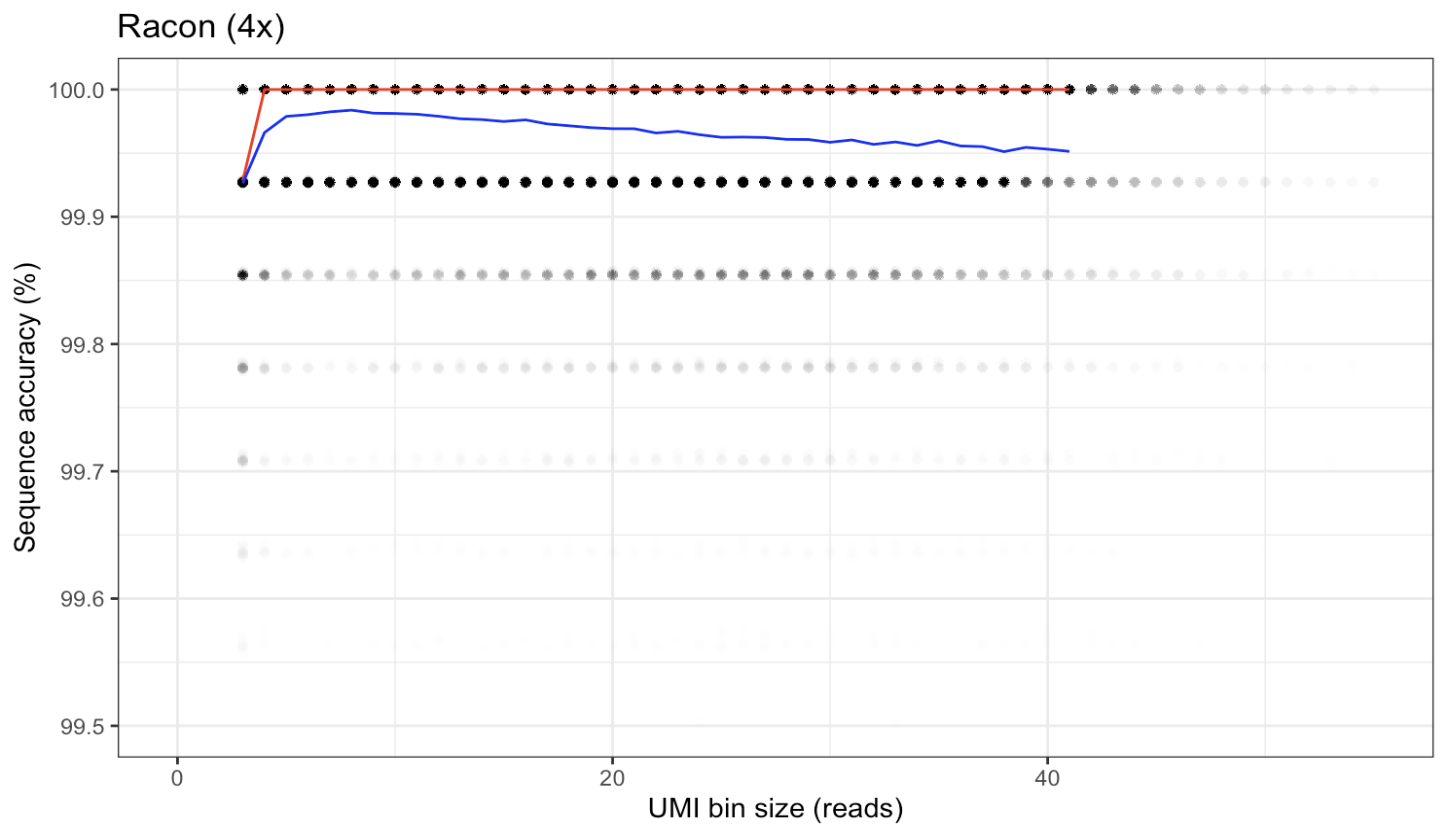

**Fig. S2.** Accuracy of UMI-based 16S rRNA gene consensus sequences (y-axis) versus UMI bin size (number of sub-reads sharing same UMI-pair; x-axis) after 4-rounds of Racon polishing, with no Medaka. The UMI-tagged 16S rRNA gene amplicons were generated from the ZymoBIOMICS Microbial Community DNA Standard. Black points represent the accuracy of individual consensus sequences, the red line represents the median accuracy of all consensus sequences at that UMI bin size, and the blue line represents the mean sequence accuracy.

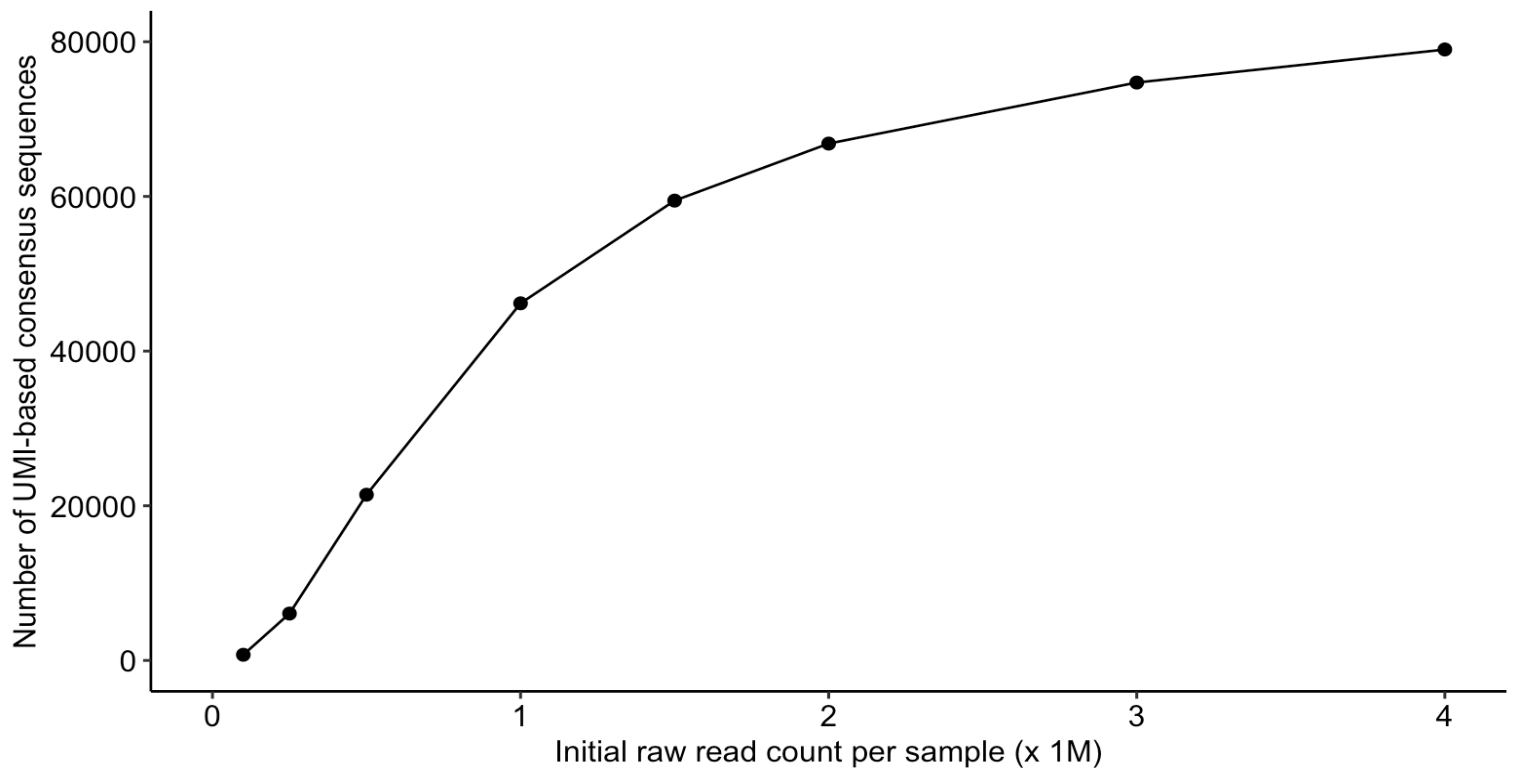

**Fig. S3.** The number of UMI-based 16S rRNA gene consensus sequences with sub-read coverage ≥3x that were recovered as a function of raw-read depth. The different read depths were achieved by randomly sub-sampling the full library pool of the ZymoBIOMICS Microbial Community DNA Standard using seqtk to specified raw-read depths, and then using those sub-sampled reads to generate UMI-based consensus sequences with the ssUMI workflow.

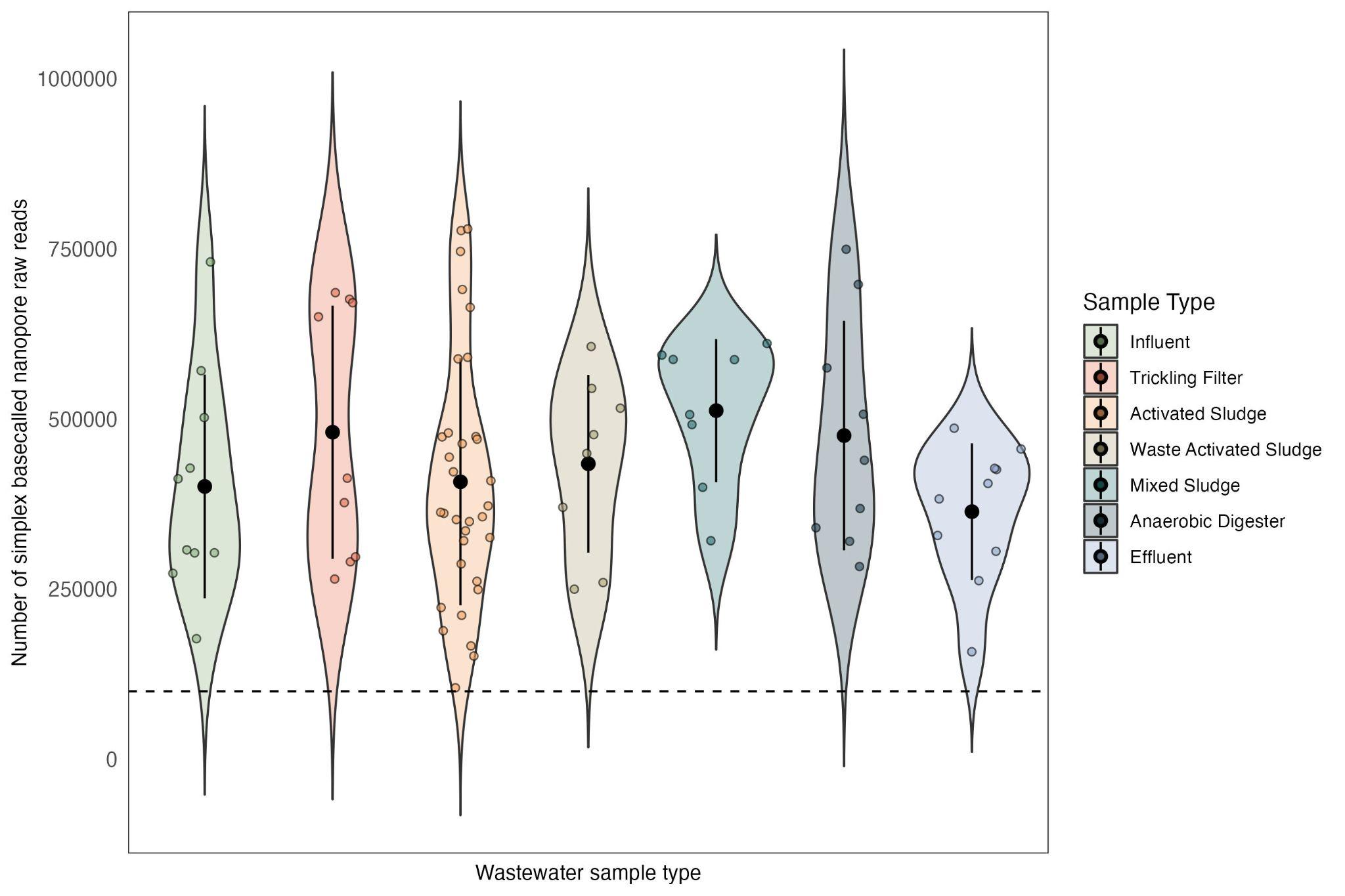

**Fig. S4.** The number of simplex Nanopore raw reads generated from different wastewater sample types. Each colored point represents one sequenced sample, the black dots represent the mean number of consensus sequences and the bars show standard deviation. The dashed line indicates 100000 raw reads.

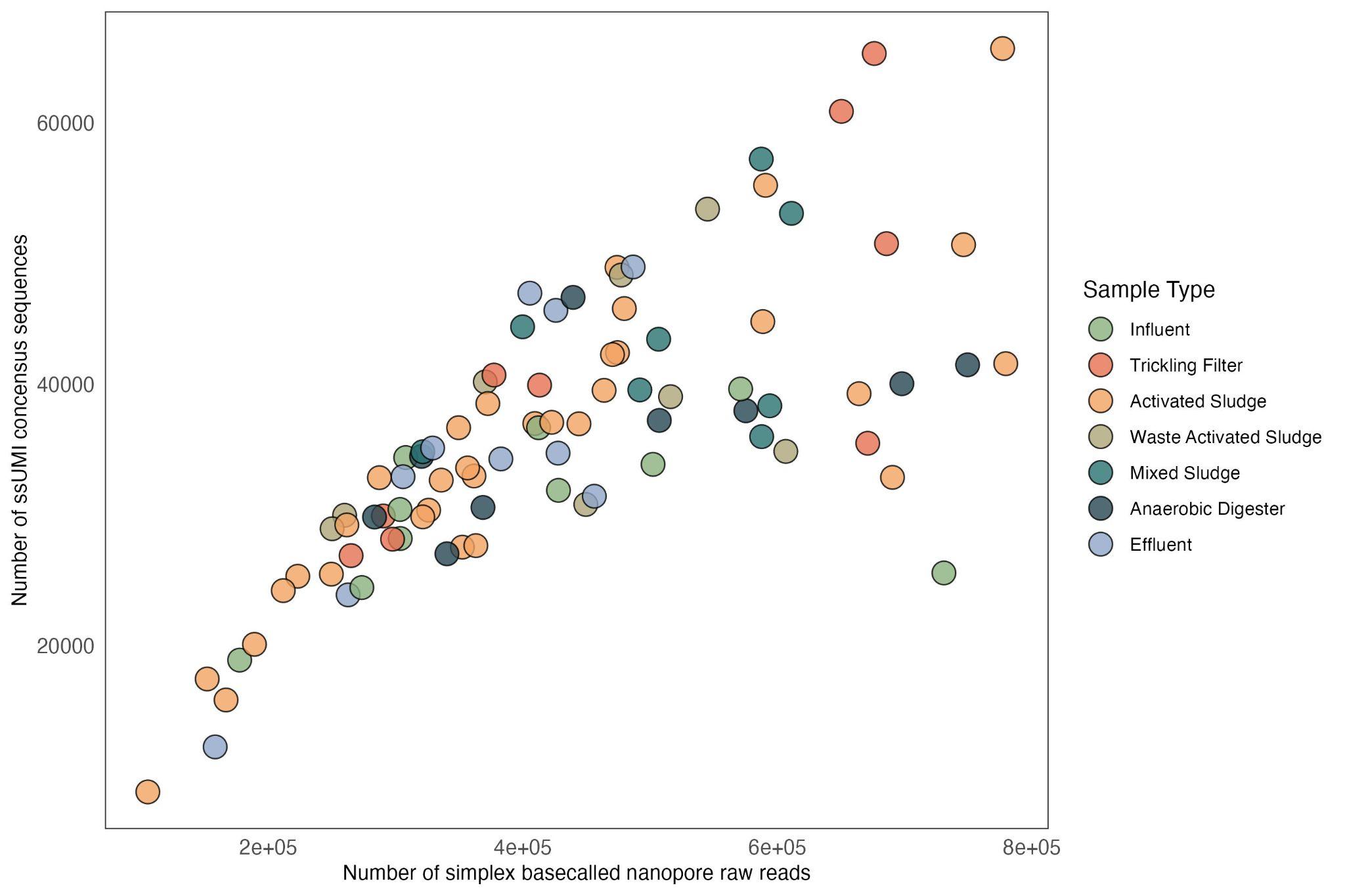

**Fig. S5.** The number of ssUMI consensus sequences generated across sequencing depths for different wastewater sample types. Each data point represents one sequenced wastewater sample, and is colored according to its sample type.

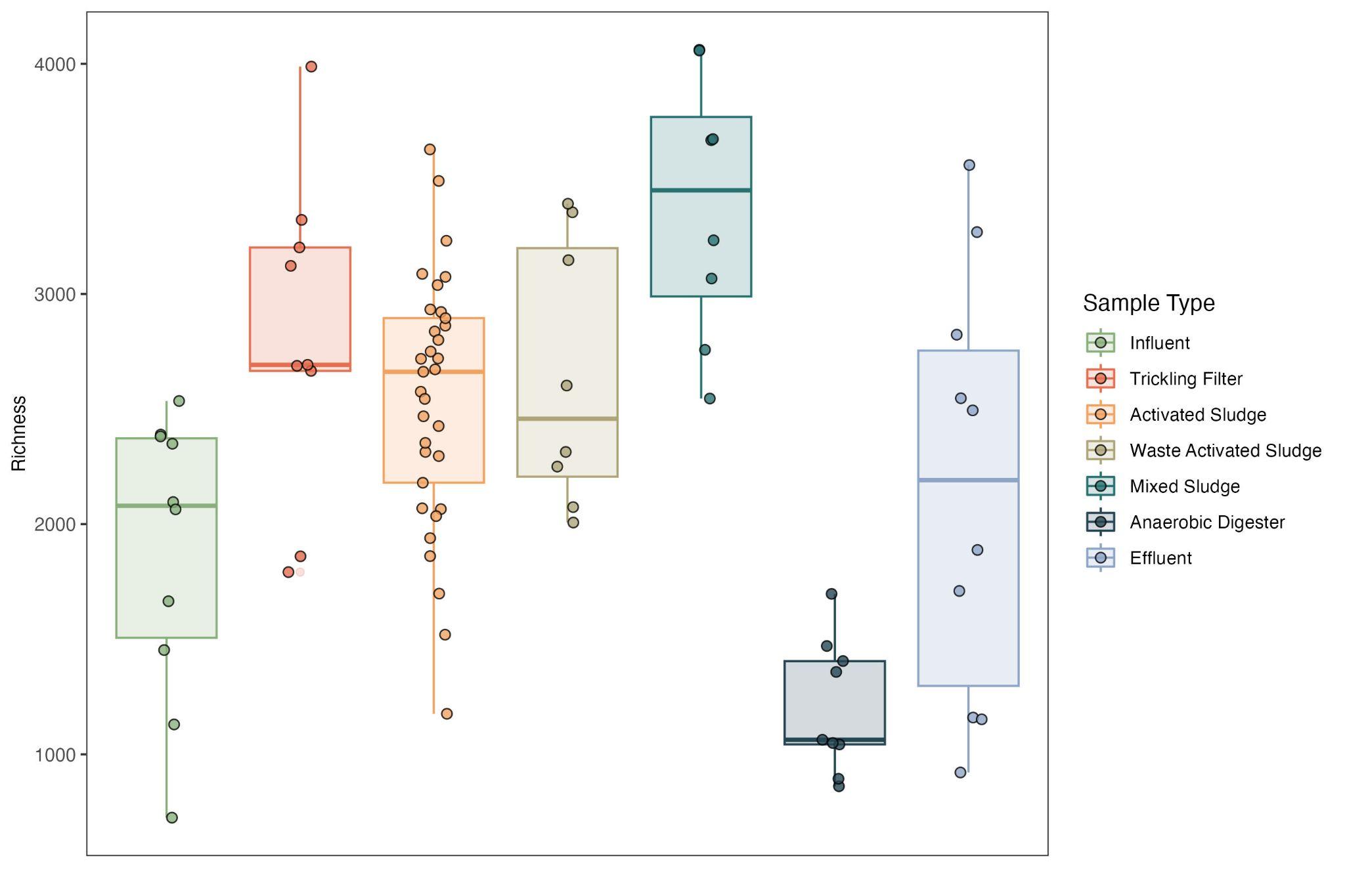

**Fig. S6.** Boxplot showing richness estimated with full-length bacterial 16S rRNA gene ASVs in various wastewater sample matrices. Each data point represents a sequenced sample, and the color indicates its sample type.

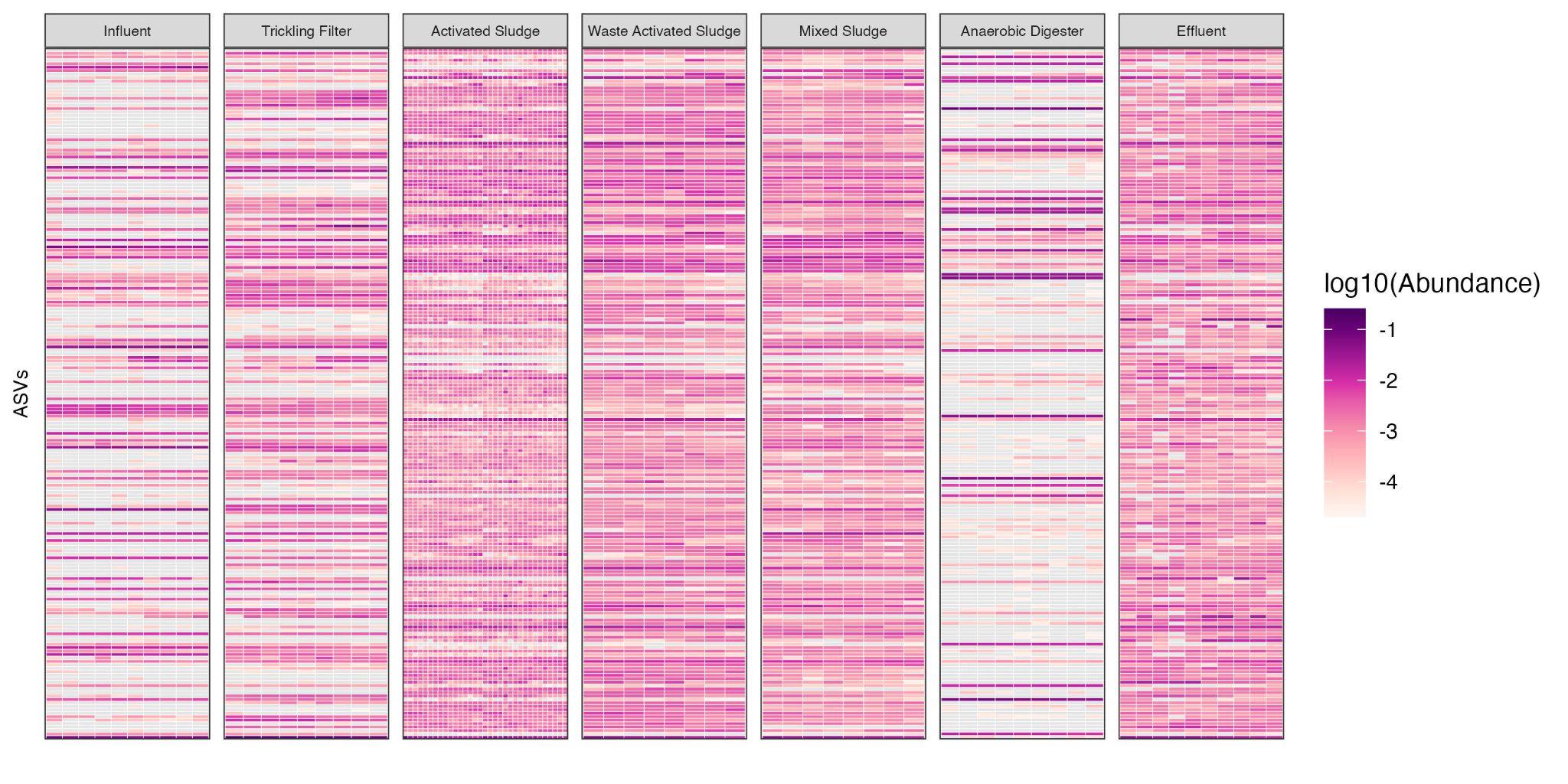

**Fig. S7.** Heatmap of bacterial ASVs generated with ssUMI consensus sequences from seven wastewater sample matrices. Only the most abundant 200 ASVs across all sample types were visualized. Zeros are shown with gray.

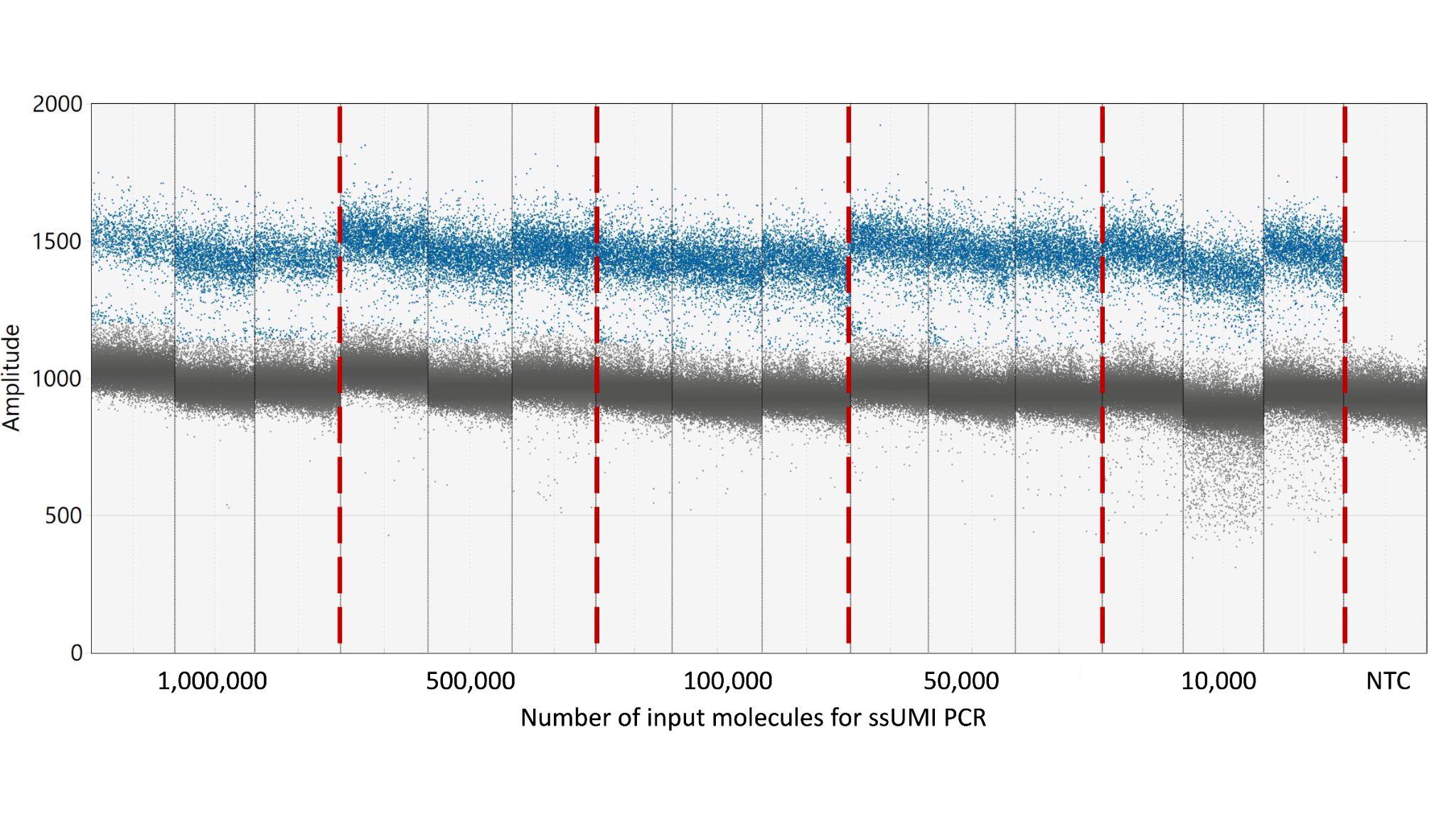

**Fig. S8.** Optimization of 16S rRNA gene copy number into the ssUMI workflow based on droplet PCR patterns of *E. coli* full-length 16S rRNA gene assay after UMI-tagging (ssUMI-PCR1), using 8F/1391R primer set and FAM-515F probe (see Supplemental Table S9). Input molecule number to ddPCR was 10000 copies per reaction. Nuclease-free was included as ddPCR no-template control (NTC). Positive (blue) and negative (grey) droplets were classified automatically using Bio-rad QX ONE Software.

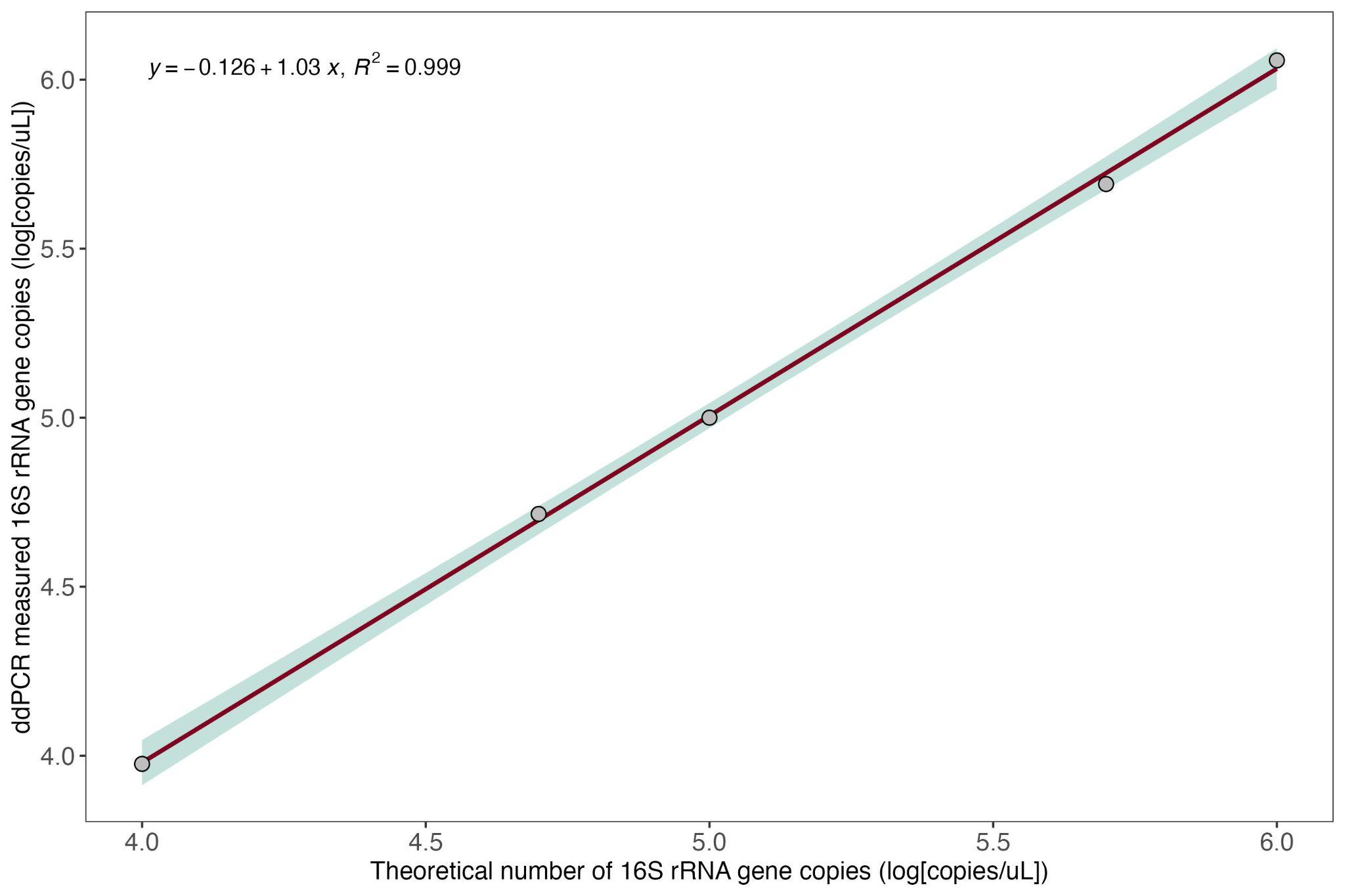

**Fig. S9.** Correlation between the ddPCR-measured (log_10_-scaled) and theoretical number (log_10_-scaled) of full-length 16S rRNA gene copies in five *E. coli* genomic DNA dilutions. The theoretical values were calculated based on an initial ddPCR quantification with ~10000 gene copies as input, before serial-dilution of the *E. coli* DNA and ddPCR quantification here.

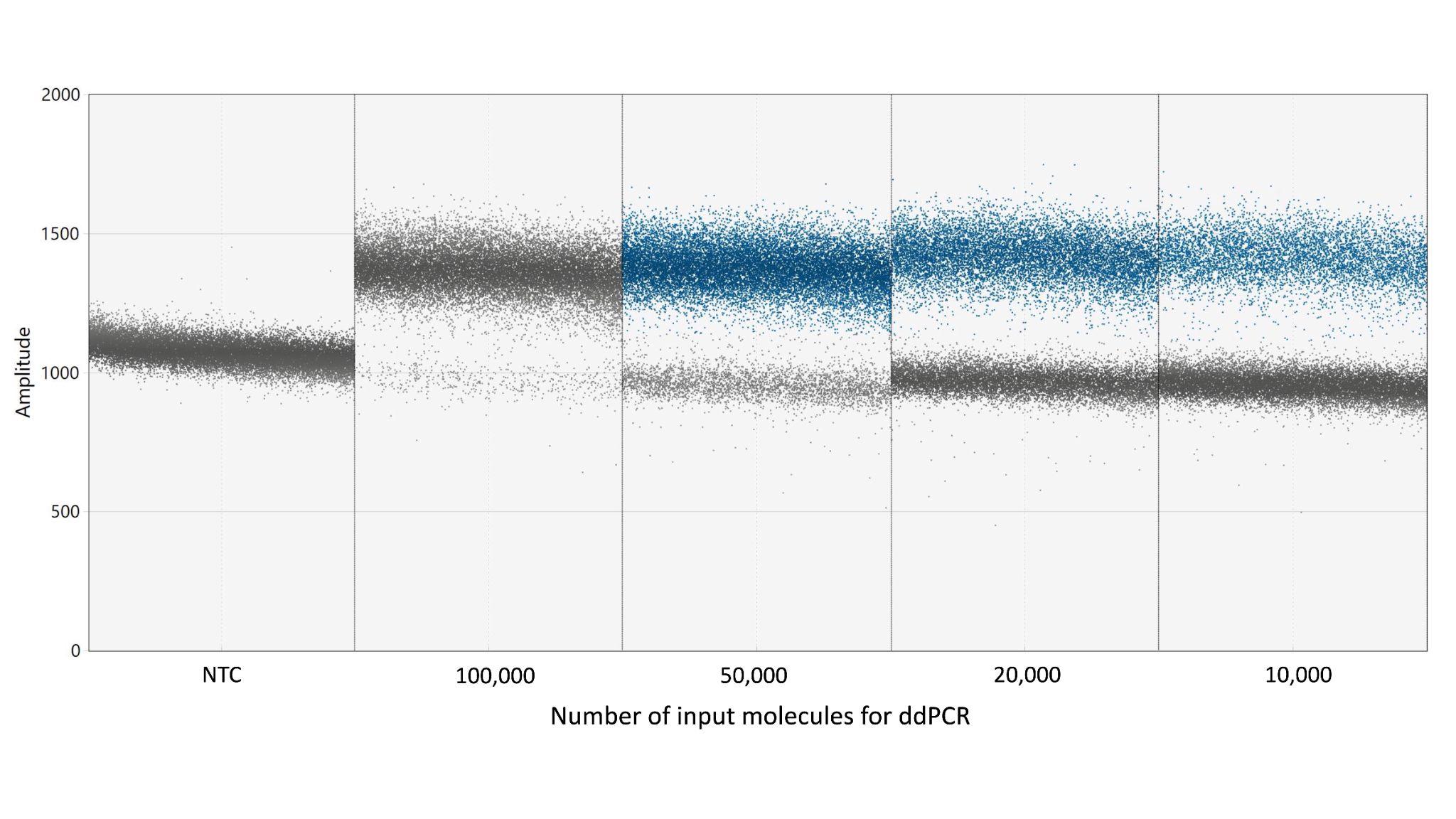

**Fig. S10.** Droplet patterns of a series-dilution of input full-length 16S rRNA gene copies into ddPCR. The ZymoBIOMICS Gut Microbiome Standard extracted with phenol-chloroform protocol was used as the DNA source. The DNA was diluted with nuclease-free water to the desired concentration, based on an initial ddPCR quantification with ~1000 gene copies as input, before ddPCR quantification here. Nuclease-free water was included as ddPCR no-template control (NTC). Positive (blue) and negative (grey) droplets were classified automatically using Bio-rad QX ONE Software. Positive droplets in the 100000 gene-copy input reaction weren't automatically classified due to the low number of negative droplets. Based on these results, an input 16S rRNA gene copy number of 50000 was chosen for the ddPCR assay going forward.

**Table S1.** Summary of computational requirements for ‘rapid’ and ‘standard’ UMI-based consensus modes, applied to process 500k UMI-tagged raw reads from the ZymoBIOMICS Microbial Community DNA Standard. The analysis was performed on a Dell EMC C4140 compute node equipped with a NVIDIA Tesla V100 (32GB) GPU and 16 CPU cores (128 Gb RAM)

| **Mode** | **CPU usage (CPU-hrs)** | **CPU maximum memory used (Gb)** | **GPU usage**  **(GPU-hrs)** | **GPU maximum memory used (Gb)** |
| --- | --- | --- | --- | --- |
| ssUMI_rapid | 2.5 | 9.8 | 0 | 0 |
| ssUMI_standard | 6.5 | 18.3 | 2.65 | 30.71 |

**Table S2.** Summary of amplicon sequence variants (ASVs) generated with ssUMI_std 16S rRNA gene consensus sequences from the six technical replicates of the ZymoBIOMICS Gut Microbiome Standard. Error counts with asterisks (*) were confirmed via BLASTn against NCBI nr (see Table S12).

| **Feature** | **Species** | **Number of errors** | **Theoretical abundance (%)** |
| --- | --- | --- | --- |
| Zotu1 | *Veillonella rogosae* | 0 | 15.87 |
| Zotu2 | *Veillonella rogosae* | 0 | 15.87 |
| Zotu3 | *Veillonella rogosae* | 0 | 15.87 |
| Zotu4 | *Faecalibacterium prausnitzii* | 0 | 17.63 |
| Zotu5 | *Faecalibacterium prausnitzii* | 0 | 17.63 |
| Zotu6 | *Faecalibacterium prausnitzii* | 0 | 17.63 |
| Zotu7 | *Bacteroides fragilis* | 0 | 9.94 |
| Zotu8 | *Bacteroides fragilis* | 0 | 9.94 |
| Zotu9 | *Bacteroides fragilis* | 0 | 9.94 |
| Zotu10 | *Bacteroides fragilis* | 0 | 9.94 |
| Zotu11 | *Faecalibacterium prausnitzii* | 0 | 17.63 |
| Zotu12 | *Faecalibacterium prausnitzii* | 0 | 17.63 |
| Zotu13 | *Bacteroides fragilis* | 0 | 9.94 |
| Zotu14 | *Bacteroides fragilis* | 0 | 9.94 |
| Zotu15 | *Bifidobacterium adolescentis* | 0 | 8.78 |
| Zotu16 | *Roseburia hominis* | 0 | 9.89 |
| Zotu17 | *Faecalibacterium prausnitzii* | 0 | 17.63 |
| Zotu18 | *Roseburia hominis* | 0 | 9.89 |
| Zotu19 | *Roseburia hominis* | 0 | 9.89 |
| Zotu20 | *Escherichia coli* | 0 | 12.12 |
| Zotu21 | *Roseburia hominis* | 0 | 9.89 |
| Zotu22 | *Prevotella corporis* | 0 | 4.98 |
| Zotu23 | *Prevotella corporis* | 0 | 4.98 |
| Zotu24 | *Fusobacterium nucleatum* | 0 | 7.49 |
| Zotu25 | *Prevotella corporis* | 0 | 4.98 |
| Zotu26 | *Prevotella corporis* | 0 | 4.98 |
| Zotu27 | *Fusobacterium nucleatum* | 0 | 7.49 |
| Zotu28 | *Fusobacterium nucleatum* | 0 | 7.49 |
| Zotu29 | *Fusobacterium nucleatum* | 0 | 7.49 |
| Zotu30 | *Fusobacterium nucleatum* | 0 | 7.49 |
| Zotu31 | *Bifidobacterium adolescentis* | 0 | 8.78 |
| Zotu32 | *Bifidobacterium adolescentis* | 0 | 8.78 |
| Zotu33 | *Escherichia coli* | 0 | 12.12 |
| Zotu34 | *Clostridioides difficille* | 0 | 2.62 |
| Zotu35 | *Escherichia coli* | 0 | 12.12 |
| Zotu36 | *Escherichia coli* | 0 | 12.12 |
| Zotu37 | *Escherichia coli* | 0 | 12.12 |
| Zotu38 | *Clostridioides difficille* | 0 | 2.62 |
| Zotu39 | *Akkermansia muciniphila* | 0 | 0.97 |
| Zotu40 | *Escherichia coli* | 0 | 12.12 |
| Zotu41 | *Lactobacillus fermentum* | 0 | 9.63 |
| Zotu42 | *Lactobacillus fermentum* | 0 | 9.63 |
| Zotu43 | *Lactobacillus fermentum* | 0 | 9.63 |
| Zotu44 | *Lactobacillus fermentum* | 0 | 9.63 |
| Zotu45 | *Lactobacillus fermentum* | 0 | 9.63 |
| Zotu46 | *Escherichia coli* | 0 | 12.12 |
| Zotu47 | *Escherichia coli* | 0* | 12.12 |
| Zotu48 | *Escherichia coli* | 0 | 12.12 |
| Zotu49 | *Escherichia coli* | 0 | 12.12 |
| Zotu50 | *Escherichia coli* | 0 | 12.12 |
| Zotu51 | *Clostridioides difficille* | 0 | 2.62 |
| Zotu52 | *Escherichia coli* | 0 | 12.12 |
| Zotu53 | *Escherichia coli* | 0* | 12.12 |
| Zotu54 | *Clostridioides difficille* | 0 | 2.62 |
| Zotu55 | *Escherichia coli* | 0* | 12.12 |
| Zotu56 | *Escherichia coli* | 0 | 12.12 |
| Zotu57 | *Escherichia coli* | 0 | 12.12 |
| Zotu58 | *Clostridioides difficille* | 0 | 2.62 |
| Zotu59 | *Clostridioides difficille* | 0 | 2.62 |
| Zotu60 | *Clostridioides difficille* | 0 | 2.62 |
| Zotu61 | *Escherichia coli* | 0 | 12.12 |
| Zotu62 | *Clostridioides difficille* | 0 | 2.62 |
| Zotu63 | *Escherichia coli* | 0 | 12.12 |
| Zotu64 | *Clostridioides difficille* | 0 | 2.62 |
| Zotu65 | *Salmonella enterica* | 0 | 0.009 |

**Table S3.** Summary of amplicon sequence variants (ASVs) generated with ssUMI_rapid 16S rRNA gene consensus sequences from the six technical replicates of the ZymoBIOMICS Gut Microbiome Standard. Error counts with asterisks (*) were confirmed via BLASTn against NCBI nr (see Table S12).

| **Feature** | **Species** | **Number of errors** | **Theoretical abundance (%)** |
| --- | --- | --- | --- |
| Zotu1 | *Veillonella rogosae* | 0 | 15.87 |
| Zotu2 | *Veillonella rogosae* | 0 | 15.87 |
| Zotu3 | *Veillonella rogosae* | 0 | 15.87 |
| Zotu4 | *Faecalibacterium prausnitzii* | 0 | 17.63 |
| Zotu5 | *Faecalibacterium prausnitzii* | 0 | 17.63 |
| Zotu6 | *Faecalibacterium prausnitzii* | 0 | 17.63 |
| Zotu7 | *Bacteroides fragilis* | 0 | 9.94 |
| Zotu8 | *Bacteroides fragilis* | 0 | 9.94 |
| Zotu9 | *Bacteroides fragilis* | 0 | 9.94 |
| Zotu10 | *Bacteroides fragilis* | 0 | 9.94 |
| Zotu11 | *Faecalibacterium prausnitzii* | 0 | 17.63 |
| Zotu12 | *Faecalibacterium prausnitzii* | 0 | 17.63 |
| Zotu13 | *Bacteroides fragilis* | 0 | 9.94 |
| Zotu14 | *Bacteroides fragilis* | 0 | 9.94 |
| Zotu15 | *Bifidobacterium adolescentis* | 0 | 8.78 |
| Zotu16 | *Faecalibacterium prausnitzii* | 0 | 17.63 |
| Zotu17 | *Roseburia hominis* | 0 | 9.89 |
| Zotu18 | *Roseburia hominis* | 0 | 9.89 |
| Zotu19 | *Escherichia coli* | 0 | 12.12 |
| Zotu20 | *Roseburia hominis* | 0 | 9.89 |
| Zotu21 | *Roseburia hominis* | 0 | 9.89 |
| Zotu22 | *Prevotella corporis* | 0 | 4.98 |
| Zotu23 | *Prevotella corporis* | 0 | 4.98 |
| Zotu24 | *Fusobacterium nucleatum* | 0 | 7.49 |
| Zotu25 | *Prevotella corporis* | 0 | 4.98 |
| Zotu26 | *Prevotella corporis* | 0 | 4.98 |
| Zotu27 | *Fusobacterium nucleatum* | 0 | 7.49 |
| Zotu28 | *Fusobacterium nucleatum* | 0 | 7.49 |
| Zotu29 | *Fusobacterium nucleatum* | 0 | 7.49 |
| Zotu30 | *Fusobacterium nucleatum* | 0 | 7.49 |
| Zotu31 | *Bifidobacterium adolescentis* | 0 | 8.78 |
| Zotu32 | *Bifidobacterium adolescentis* | 0 | 8.78 |
| Zotu33 | *Clostridioides difficille* | 0 | 2.62 |
| Zotu34 | *Escherichia coli* | 0 | 12.12 |
| Zotu35 | *Escherichia coli* | 0 | 12.12 |
| Zotu36 | *Escherichia coli* | 0 | 12.12 |
| Zotu37 | *Escherichia coli* | 0 | 12.12 |
| Zotu38 | *Akkermansia muciniphila* | 0 | 0.97 |
| Zotu39 | *Clostridioides difficille* | 0 | 2.62 |
| Zotu40 | *Escherichia coli* | 0 | 12.12 |
| Zotu41 | *Lactobacillus fermentum* | 0 | 9.63 |
| Zotu42 | *Lactobacillus fermentum* | 0 | 9.63 |
| Zotu43 | *Lactobacillus fermentum* | 0 | 9.63 |
| Zotu44 | *Lactobacillus fermentum* | 0 | 9.63 |
| Zotu45 | *Lactobacillus fermentum* | 0 | 9.63 |
| Zotu46 | *Escherichia coli* | 0 | 12.12 |
| Zotu47 | *Escherichia coli* | 0* | 12.12 |
| Zotu48 | *Escherichia coli* | 0 | 12.12 |
| Zotu49 | *Escherichia coli* | 0 | 12.12 |
| Zotu50 | *Escherichia coli* | 0 | 12.12 |
| Zotu51 | *Clostridioides difficille* | 0 | 2.62 |
| Zotu52 | *Clostridioides difficille* | 0 | 2.62 |
| Zotu53 | *Escherichia coli* | 0 | 12.12 |
| Zotu54 | *Escherichia coli* | 0* | 12.12 |
| Zotu55 | *Escherichia coli* | 0 | 12.12 |
| Zotu56 | *Escherichia coli* | 0* | 12.12 |
| Zotu57 | *Clostridioides difficille* | 0 | 2.62 |
| Zotu58 | *Escherichia coli* | 0 | 12.12 |
| Zotu59 | *Clostridioides difficille* | 0 | 2.62 |
| Zotu60 | *Clostridioides difficille* | 0 | 2.62 |
| Zotu61 | *Clostridioides difficille* | 0 | 2.62 |
| Zotu62 | *Escherichia coli* | 0 | 12.12 |
| Zotu63 | *Escherichia coli* | 0 | 12.12 |
| Zotu64 | *Clostridioides difficille* | 0 | 2.62 |

**Table S4.** Summary of amplicon sequence variants (ASVs) generated with quality-filtered 16S rRNA gene amplicons from the six technical replicates of the ZymoBIOMICS Gut Microbiome Standard.

| **Feature** | **Species** | **Number of errors** | **Theoretical abundance (%)** |
| --- | --- | --- | --- |
| Zotu1 | *Veillonella rogosae* | 0 | 15.87 |
| Zotu2 | *Faecalibacterium prausnitzii* | 0 | 17.63 |
| Zotu3 | *Veillonella rogosae* | 0 | 15.87 |
| Zotu4 | *Veillonella rogosae* | 0 | 15.87 |
| Zotu5 | *Faecalibacterium prausnitzii* | 0 | 17.63 |
| Zotu6 | *Bacteroides fragilis* | 0 | 9.94 |
| Zotu7 | *Faecalibacterium prausnitzii* | 0 | 17.63 |
| Zotu8 | *Bacteroides fragilis* | 0 | 9.94 |
| Zotu9 | *Faecalibacterium prausnitzii* | 0 | 17.63 |
| Zotu10 | *Bacteroides fragilis* | 0 | 9.94 |
| Zotu11 | *Bacteroides fragilis* | 0 | 9.94 |
| Zotu12 | *Faecalibacterium prausnitzii* | 0 | 17.63 |
| Zotu13 | *Bacteroides fragilis* | 0 | 9.94 |
| Zotu14 | *Bacteroides fragilis* | 0 | 9.94 |
| Zotu15 | *Fusobacterium nucleatum* | 0 | 7.49 |
| Zotu16 | *Faecalibacterium prausnitzii* | 0 | 17.63 |
| Zotu17 | *Escherichia coli* | 0 | 12.12 |
| Zotu18 | *Bifidobacterium adolescentis* | 0 | 8.78 |
| Zotu19 | *Prevotella corporis* | 0 | 4.98 |
| Zotu20 | *Prevotella corporis* | 0 | 4.98 |
| Zotu21 | *Fusobacterium nucleatum* | 0 | 7.49 |
| Zotu22 | *Fusobacterium nucleatum* | 0 | 7.49 |
| Zotu23 | *Fusobacterium nucleatum* | 0 | 7.49 |
| Zotu24 | *Roseburia hominis* | 0 | 9.89 |
| Zotu25 | *Fusobacterium nucleatum* | 0 | 7.49 |
| Zotu26 | *Prevotella corporis* | 0 | 4.98 |
| Zotu27 | *Roseburia hominis* | 0 | 9.89 |
| Zotu28 | *Roseburia hominis* | 0 | 9.89 |
| Zotu29 | *Prevotella corporis* | 0 | 4.98 |
| Zotu30 | *Roseburia hominis* | 0 | 9.89 |
| Zotu31 | *Lactobacillus fermentum* | 0 | 9.63 |
| Zotu32 | *Lactobacillus fermentum* | 0 | 9.63 |
| Zotu33 | *Lactobacillus fermentum* | 0 | 9.63 |
| Zotu34 | *Lactobacillus fermentum* | 0 | 9.63 |
| Zotu35 | *Clostridioides difficille* | 0 | 2.62 |
| Zotu36 | *Escherichia coli* | 0 | 12.12 |
| Zotu37 | *Lactobacillus fermentum* | 0 | 9.63 |
| Zotu38 | *Escherichia coli* | 0 | 12.12 |
| Zotu39 | *Escherichia coli* | 0 | 12.12 |
| Zotu40 | *Clostridioides difficille* | 0 | 2.62 |
| Zotu41 | *Bifidobacterium adolescentis* | 0 | 8.78 |
| Zotu42 | *Bifidobacterium adolescentis* | 0 | 8.78 |
| Zotu43 | *Akkermansia muciniphila* | 0 | 0.97 |
| Zotu44 | *Faecalibacterium prausnitzii* | 2 | 17.63 |
| Zotu45 | *Escherichia coli* | 0 | 12.12 |
| Zotu46 | *Escherichia coli* | 0 | 12.12 |
| Zotu47 | *Escherichia coli* | 0 | 12.12 |
| Zotu48 | *Escherichia coli* | 0 | 12.12 |
| Zotu49 | *Faecalibacterium prausnitzii* | 0 | 17.63 |
| Zotu50 | *Clostridioides difficille* | 0 | 2.62 |
| Zotu51 | *Escherichia coli* | 0 | 12.12 |
| Zotu52 | *Escherichia coli* | 0 | 12.12 |
| Zotu53 | *Escherichia coli* | 0 | 12.12 |
| Zotu54 | *Clostridioides difficille* | 0 | 2.62 |
| Zotu55 | *Clostridioides difficille* | 0 | 2.62 |
| Zotu56 | *Clostridioides difficille* | 0 | 2.62 |
| Zotu57 | *Clostridioides difficille* | 0 | 2.62 |
| Zotu58 | *Escherichia coli* | 0 | 12.12 |
| Zotu59 | *Clostridioides difficille* | 0 | 2.62 |
| Zotu60 | *Escherichia coli* | 0 | 12.12 |
| Zotu61 | *Clostridioides difficille* | 0 | 2.62 |
| Zotu62 | *Escherichia coli* | 0 | 12.12 |
| Zotu63 | *Faecalibacterium prausnitzii* | 2 | 17.63 |
| Zotu64 | *Faecalibacterium prausnitzii* | 0 | 17.63 |
| Zotu65 | *Escherichia coli* | 0 | 12.12 |
| Zotu66 | *Escherichia coli* | 0 | 12.12 |
| Zotu67 | *Escherichia coli* | 0 | 12.12 |
| Zotu68 | *Faecalibacterium prausnitzii* | 2 | 17.63 |
| Zotu69 | *Veillonella rogosae* | 1 | 15.87 |
| Zotu70 | *Faecalibacterium prausnitzii* | 2 | 17.63 |

**Table S5.** Summary of operational taxonomic units (OTUs) with 97% identity threshold, generated with ssUMI_std 16S rRNA gene consensus sequences from the six technical replicates of the ZymoBIOMICS Gut Microbiome Standard.

| **Feature** | **Species** | **Number of errors** | **Theoretical abundance (%)** |
| --- | --- | --- | --- |
| Otu1 | *Veillonella rogosae* | 0 | 15.87 |
| Otu2 | *Faecalibacterium prausnitzii* | 0 | 17.63 |
| Otu3 | *Bacteroides fragilis* | 0 | 9.94 |
| Otu4 | *Bifidobacterium adolescentis* | 0 | 8.78 |
| Otu5 | *Roseburia hominis* | 0 | 9.89 |
| Otu6 | *Escherichia coli* | 0 | 12.12 |
| Otu7 | *Prevotella corporis* | 0 | 4.98 |
| Otu8 | *Fusobacterium nucleatum* | 0 | 7.49 |
| Otu9 | *Clostridioides difficille* | 0 | 2.62 |
| Otu10 | *Akkermansia muciniphila* | 0 | 0.97 |
| Otu11 | *Lactobacillus fermentum* | 0 | 9.63 |
| Otu12 | *Salmonella enterica* | 0 | 0.009 |
| Otu14 | *Enterococcus faecalis* | 0 | 9.00E-04 |

**Table S6.** Summary of operational taxonomic units (OTUs) with 97% identity threshold, generated with ssUMI_rapid 16S rRNA gene consensus sequences from the six technical replicates of the ZymoBIOMICS Gut Microbiome Standard.

| **Feature** | **Species** | **Number of errors** | **Theoretical abundance (%)** |
| --- | --- | --- | --- |
| Otu1 | *Veillonella rogosae* | 0 | 15.87 |
| Otu2 | *Faecalibacterium prausnitzii* | 0 | 17.63 |
| Otu3 | *Bacteroides fragilis* | 0 | 9.94 |
| Otu4 | *Bifidobacterium adolescentis* | 0 | 8.78 |
| Otu5 | *Roseburia hominis* | 0 | 9.89 |
| Otu6 | *Escherichia coli* | 0 | 12.12 |
| Otu7 | *Prevotella corporis* | 0 | 4.98 |
| Otu8 | *Fusobacterium nucleatum* | 0 | 7.49 |
| Otu9 | *Clostridioides difficille* | 0 | 2.62 |
| Otu10 | *Akkermansia muciniphila* | 0 | 0.97 |
| Otu11 | *Lactobacillus fermentum* | 0 | 9.63 |
| Otu13 | *Salmonella enterica* | 0 | 0.009 |
| Otu14 | *Enterococcus faecalis* | 0 | 9.00E-04 |

**Table S7.** Summary of operational taxonomic units (OTUs) with 97% identity threshold, generated with quality-filtered Nanopore simples read 16S rRNA gene amplicons from the six technical replicates of the ZymoBIOMICS Gut Microbiome Standard.

| **Feature** | **Species** | **Number of errors** | **Theoretical abundance (%)** |
| --- | --- | --- | --- |
| Otu1 | *Veillonella rogosae* | 0 | 15.87 |
| Otu2 | *Faecalibacterium prausnitzii* | 0 | 17.63 |
| Otu3 | *Bacteroides fragilis* | 0 | 9.94 |
| Otu4 | *Fusobacterium nucleatum* | 0 | 7.49 |
| Otu5 | *Bifidobacterium adolescentis* | 0 | 8.78 |
| Otu6 | *Prevotella corporis* | 0 | 4.98 |
| Otu7 | *Escherichia coli* | 0 | 12.12 |
| Otu8 | *Roseburia hominis* | 0 | 9.89 |
| Otu9 | *Lactobacillus fermentum* | 0 | 9.63 |
| Otu10 | *Clostridioides difficille* | 0 | 2.62 |
| Otu11 | *Akkermansia muciniphila* | 0 | 0.97 |
| Otu12 | *Salmonella enterica* | 0 | 0.009 |

**Table S8.** Summary of ssUMI amplicon concentrations, sequencing throughputs and ssUMI consensus sequences processed with ssUMI_std mode of wastewater samples.

| Sample ID | Date Sampled | Sample Type | ssUMI amplicon concentration (ng/uL) | Raw reads | ssUMI consensus sequences | Accession |
| --- | --- | --- | --- | --- | --- | --- |
| Inf_06/21/2022 | 6/21/2022 | Influent | 16.1 | 307,810 | 34,403 | SAMN35219784 |
| Eff_06/21/2022 | 6/21/2022 | Effluent | 15.6 | 425,790 | 45,670 | SAMN35219785 |
| TF_06/21/2022 | 6/22/2022 | Trickling Filter | 9.22 | 290,247 | 29,935 | SAMN35219786 |
| AS-1_06/22/2022 | 6/22/2022 | Activated Sludge | 11.3 | 349,366 | 36,691 | SAMN35219787 |
| AS-2_06/22/2022 | 6/22/2022 | Activated Sludge | 10.4 | 105,217 | 8,812 | SAMN35219788 |
| WAS_06/22/2022 | 6/22/2022 | Waste Activated Sludge | 6.38 | 259,793 | 29,987 | SAMN35219789 |
| AD_06/22/2022 | 6/22/2022 | Anaerobic Digester | 7.81 | 320,304 | 34,495 | SAMN35219790 |
| Inf_06/27/2022 | 6/27/2022 | Influent | 7.8 | 427,841 | 31,895 | SAMN35219791 |
| Eff_06/27/2022 | 6/27/2022 | Effluent | 16.7 | 405,366 | 46,986 | SAMN35219792 |
| TF_06/27/2022 | 6/27/2022 | Trickling Filter | 16.7 | 685,584 | 50,777 | SAMN35219793 |
| AS-1_06/28/2022 | 6/28/2022 | Activated Sludge | 11.3 | 463,694 | 39,550 | SAMN35219794 |
| AS-2_06/28/2022 | 6/28/2022 | Activated Sludge | 17.2 | 287,151 | 32,865 | SAMN35219795 |
| WAS_06/28/2022 | 6/28/2022 | Waste Activated Sludge | 13.5 | 370,334 | 40,193 | SAMN35219796 |
| TSS_06/28/2022 | 6/28/2022 | Mixed Sludge | 19.4 | 399,623 | 44,411 | SAMN35219797 |
| AD_06/28/2022 | 6/28/2022 | Anaerobic Digester | 6.86 | 368,525 | 30,585 | SAMN35219798 |
| AS-1_06/30/2022 | 6/30/2022 | Activated Sludge | 10.9 | 325,837 | 30,377 | SAMN35219799 |
| AS-2_06/30/2022 | 6/30/2022 | Activated Sludge | 13.9 | 473,827 | 48,960 | SAMN35219800 |
| Inf_07/04/2022 | 7/4/2022 | Influent | 8.19 | 303,519 | 28,208 | SAMN35219801 |
| Eff_07/04/2022 | 7/4/2022 | Effluent | 17.1 | 305,711 | 32,940 | SAMN35219802 |
| TF_07/04/2022 | 7/4/2022 | Trickling Filter | 8.01 | 264,899 | 26,899 | SAMN35219803 |
| AS-1_07/05/2022 | 7/5/2022 | Activated Sludge | 9.42 | 223,020 | 25,319 | SAMN35219804 |
| AS-2_07/05/2022 | 7/5/2022 | Activated Sludge | 6.96 | 249,430 | 25,480 | SAMN35219805 |
| WAS_07/05/2022 | 7/5/2022 | Waste Activated Sludge | 12.8 | 250,006 | 28,954 | SAMN35219806 |
| TSS_07/05/2022 | 7/5/2022 | Mixed Sludge | 16.8 | 321,237 | 34,851 | SAMN35219807 |
| AD_07/05/2022 | 7/5/2022 | Anaerobic Digester | 22.5 | 439,339 | 46,665 | SAMN35219808 |
| AS-1_07/07/2022 | 7/7/2022 | Activated Sludge | 11.3 | 361,467 | 32,994 | SAMN35219809 |
| AS-2_07/07/2022 | 7/7/2022 | Activated Sludge | 13 | 261,550 | 29,247 | SAMN35219810 |
| Inf_07/11/2022 | 7/11/2022 | Influent | 8.09 | 177,331 | 18,901 | SAMN35219811 |
| Eff_07/11/2022 | 7/11/2022 | Effluent | 24.4 | 158,119 | 12,258 | SAMN35219812 |
| TF_07/11/2022 | 7/11/2022 | Trickling Filter | 19.3 | 377,167 | 40,718 | SAMN35219813 |
| AS-1_07/12/2022 | 7/12/2022 | Activated Sludge | 17 | 211,611 | 24,221 | SAMN35219814 |
| AS-2_07/12/2022 | 7/12/2022 | Activated Sludge | 20.7 | 166,670 | 15,850 | SAMN35219815 |
| WAS_07/12/2022 | 7/12/2022 | Waste Activated Sludge | 5.74 | 449,332 | 30,814 | SAMN35219816 |
| TSS_07/12/2022 | 7/12/2022 | Mixed Sludge | 7.48 | 593,870 | 38,365 | SAMN35219817 |
| AD_07/12/2022 | 7/12/2022 | Anaerobic Digester | 10.9 | 507,055 | 37,247 | SAMN35219818 |
| AS-1_07/14/2022 | 7/14/2022 | Activated Sludge | 15.4 | 746,100 | 50,695 | SAMN35219819 |
| AS-2_07/14/2022 | 7/14/2022 | Activated Sludge | 3.67 | 151,834 | 17,457 | SAMN35219820 |
| Inf_07/18/2022 | 7/18/2022 | Influent | 6.68 | 730,677 | 25,572 | SAMN35219821 |
| Eff_07/18/2022 | 7/18/2022 | Effluent | 10 | 328,973 | 35,135 | SAMN35219822 |
| TF_07/18/2022 | 7/18/2022 | Trickling Filter | 19.6 | 650,147 | 60,902 | SAMN35219823 |
| AS-1_07/19/2022 | 7/19/2022 | Activated Sludge | 12.6 | 372,347 | 38,538 | SAMN35219824 |
| AS-2_07/19/2022 | 7/19/2022 | Activated Sludge | 9.54 | 779,276 | 41,592 | SAMN35219825 |
| WAS_07/19/2022 | 7/19/2022 | Waste Activated Sludge | 9.78 | 606,447 | 34,876 | SAMN35219826 |
| TSS_07/19/2022 | 7/19/2022 | Mixed Sludge | 9.57 | 587,467 | 36,005 | SAMN35219827 |
| AD_07/19/2022 | 7/19/2022 | Anaerobic Digester | 9.28 | 697,629 | 40,056 | SAMN35219828 |
| AS-1_07/21/2022 | 7/21/2022 | Activated Sludge | 12.6 | 588,398 | 44,812 | SAMN35219829 |
| AS-2_07/21/2022 | 7/21/2022 | Activated Sludge | 7.41 | 470,281 | 42,290 | SAMN35219830 |
| Inf_07/25/2022 | 7/25/2022 | Influent | 7.25 | 502,146 | 33,884 | SAMN35219831 |
| Eff_07/25/2022 | 7/25/2022 | Effluent | 5.7 | 456,011 | 31,444 | SAMN35219832 |
| TF_07/25/2022 | 7/25/2022 | Trickling Filter | 5.3 | 732 | n/a | n/a |
| AS-1_07/26/2022 | 7/26/2022 | Activated Sludge | 2.77 | 1,255 | n/a | n/a |
| AS-2_07/26/2022 | 7/26/2022 | Activated Sludge | 7.96 | 352,334 | 27,547 | SAMN35219833 |
| WAS_07/26/2022 | 7/26/2022 | Waste Activated Sludge | 8.38 | 154 | n/a | n/a |
| TSS_07/26/2022 | 7/26/2022 | Mixed Sludge | 12 | 506,530 | 43,462 | SAMN35219834 |
| AD_07/26/2022 | 7/26/2022 | Anaerobic Digester | 9.97 | 575,072 | 37,986 | SAMN35219835 |
| AS-1_07/28/2022 | 7/28/2022 | Activated Sludge | 24.9 | 409,258 | 37,008 | SAMN35219836 |
| AS-2_07/28/2022 | 7/28/2022 | Activated Sludge | 20.7 | 362,929 | 27,664 | SAMN35219837 |
| Inf_08/01/2022 | 8/1/2022 | Influent | 14.1 | 303,335 | 30,432 | SAMN35219838 |
| Eff_08/01/2022 | 8/1/2022 | Effluent | 6.75 | 262,585 | 23,894 | SAMN35219839 |
| TF_08/01/2022 | 8/1/2022 | Trickling Filter | 8.79 | 297,476 | 28,156 | SAMN35219840 |
| AS-1_08/02/2022 | 8/2/2022 | Activated Sludge | 18.9 | 479,585 | 45,808 | SAMN35219841 |
| AS-2_08/02/2022 | 8/2/2022 | Activated Sludge | 19.2 | 335,738 | 32,677 | SAMN35219842 |
| WAS_08/02/2022 | 8/2/2022 | Waste Activated Sludge | 28.8 | 515,916 | 39,065 | SAMN35219843 |
| TSS_08/02/2022 | 8/2/2022 | Mixed Sludge | 11.4 | 610,911 | 53,096 | SAMN35219844 |
| AD_08/02/2022 | 8/2/2022 | Anaerobic Digester | 2.14 | 340,222 | 27,037 | SAMN35219845 |
| AS-1_08/04/2022 | 8/4/2022 | Activated Sludge | 9.89 | 590,533 | 55,241 | SAMN35219846 |
| AS-2_08/04/2022 | 8/4/2022 | Activated Sludge | 13.2 | 443,947 | 36,988 | SAMN35219847 |
| Inf_08/08/2022 | 8/8/2022 | Influent | 7.79 | 412,199 | 36,687 | SAMN35219848 |
| Eff_08/08/2022 | 8/8/2022 | Effluent | 6.65 | 427,528 | 34,742 | SAMN35219849 |
| TF_08/08/2022 | 8/8/2022 | Trickling Filter | 9.57 | 413,012 | 39,947 | SAMN35219850 |
| AS-1_08/09/2022 | 8/9/2022 | Activated Sludge | 10.8 | 422,529 | 37,099 | SAMN35219851 |
| AS-2_08/09/2022 | 8/9/2022 | Activated Sludge | 9.07 | 663,991 | 39,294 | SAMN35219852 |
| WAS_08/09/2022 | 8/9/2022 | Waste Activated Sludge | 15 | 477,113 | 48,374 | SAMN35219853 |
| TSS_08/09/2022 | 8/9/2022 | Mixed Sludge | 9.13 | 491,750 | 39,581 | SAMN35219854 |
| AD_08/09/2022 | 8/9/2022 | Anaerobic Digester | 12.9 | 283,337 | 29,851 | SAMN35219855 |
| AS-1_08/11/2022 | 8/11/2022 | Activated Sludge | 7.07 | 188,980 | 20,108 | SAMN35219856 |
| AS-2_08/11/2022 | 8/11/2022 | Activated Sludge | 10.3 | 356,471 | 33,611 | SAMN35219857 |
| Inf_08/15/2022 | 8/15/2022 | Influent | 20.2 | 273,345 | 24,453 | SAMN35219858 |
| Eff_08/15/2022 | 8/15/2022 | Effluent | 15.7 | 486,500 | 48,993 | SAMN35219859 |
| TF_08/15/2022 | 8/15/2022 | Trickling Filter | 11.8 | 675,947 | 65,331 | SAMN35219860 |
| AS-1_08/16/2022 | 8/16/2022 | Activated Sludge | 4.45 | 776,782 | 65,711 | SAMN35219861 |
| AS-2_08/16/2022 | 8/16/2022 | Activated Sludge | 4.44 | 690,281 | 32,888 | SAMN35219862 |
| WAS_08/16/2022 | 8/16/2022 | Waste Activated Sludge | 13.3 | 544,966 | 53,421 | SAMN35219863 |
| TSS_08/16/2022 | 8/16/2022 | Mixed Sludge | 15.9 | 587,174 | 57,251 | SAMN35219864 |
| AD_08/16/2022 | 8/16/2022 | Anaerobic Digester | 5.63 | 749,280 | 41,502 | SAMN35219865 |
| AS-1_08/18/2022 | 8/18/2022 | Activated Sludge | 6.68 | 321,210 | 29,863 | SAMN35219866 |
| AS-2_08/18/2022 | 8/18/2022 | Activated Sludge | 11.2 | 474,192 | 42,436 | SAMN35219867 |
| Inf_08/22/2022 | 8/22/2022 | Influent | 10 | 570,971 | 39,653 | SAMN35219868 |
| Eff_08/22/2022 | 8/22/2022 | Effluent | 9.13 | 382,601 | 34,298 | SAMN35219869 |
| TF_08/22/2022 | 8/22/2022 | Trickling Filter | 5.86 | 670,827 | 35,498 | SAMN35219870 |

**Table S9.** Per-sample cost estimates for 16S rRNA gene amplicon sequencing with either the ssUMI workflow and sequencing on ONT (V1-V9), with PacBio HiFi sequencing, or with Illumina (V4-V5). The per-sample cost was estimated based on a target per-sample throughput of ~40000 ssUMI consensus sequences (ONT), ~40000 HiFi sequences (PacBio), ~10000 LoopSeq synthetic long-reads, and ~50000 raw-reads (Illumina V4-V5). All prices were shown in USD and rounded to the nearest integer. Local taxes and shipping were included in estimates. These costs generally assume that a user has an ONT PromethION in their lab, but would have to contract sequencing facilities for PacBio and Illumina sequencing.

| **Item** | **V1-V9 with ssUMI on ONT PromethION** | **V1-V9 with HiFi on PacBio Sequel II system** | **V1-V9 synthetic long-reads with LoopSeq on Illumina NextSeq** | **V4-V5 short-reads on Illumina MiSeq** |
| --- | --- | --- | --- | --- |
| ddPCR quantification | $4 | NA | NA | NA |
| 16S rRNA gene amplicon generation | $16**^a^** | $5^a^ | NA | $5^a^ |
| Sequencing library preparation and QC | $5**^a^** | $8^c^ | $46^e^ | $2^a^ |
| Sequencing | $9^b^ | $23^d^ | $38^f^ | $5^h^ |
| **Total price per sample** | $34 | $36 | $84^g^ | $10 |

^a^ Based on costs incurred for this study.

^b^ Cost for an ONT R.10.4.1 flow-cell capable of generating 100 Gbp, and assuming 96 samples are multiplexed. This would generate ~40000 ssUMI sequences per sample.

^c^ Cost for HiFi library preparation at the University of Washington PacBio Sequencing Services^2^, based on 96 samples.

^d^ Cost for single SMRT 8M cell on PacBio Sequel II at the University of Washington PacBio Sequencing Services^3^, assuming 96 samples are multiplexed. This would generate ~3.5M HiFi 16S rRNA gene amplicon reads^4^, or ~40,000 reads per sample.

^e^ Cost of library preparation based on 96 barcoded samples using LoopSeq Kit for 16S rRNA on Illumina, generating ~10000 UMI-tagged molecules per sample.

^f^ Based on a price of $5000 USD per NextSeq 2000 P2 flow-cell in 2x150 bp mode, generating 400 M paired-end reads, and providing >30x coverage for ~10000 full-length 16S rRNA gene molecules in 130 samples.

^g^ It should be noted that this price would likely be lower if the Element Biosciences AVITI platform was used instead of Illumina.

^h^ Based on a price of $2000 USD per MiSeq Reagent Kit v3 (600 cycle), running 395 samples + 1 negative control. This would generate ~ 20 M merged reads, or ~50000 reads/sample.

**Table S10.** Sequences of primer and probes used in full-length 16S rRNA gene ddPCR and ssUMI-PCR.

| **Primer/Probe** | **Primer purification** | **Primer sequence 5’ to 3’** | **Reference** |
| --- | --- | --- | --- |
| **Full-length 16S rRNA gene ddPCR Assay** | | | |
| ddPCR-8F | Standard Desalting | AGRGTTYGATYMTGGCTCAG | Callahan et. al., 2019 ^5^ |
| ddPCR-1391R | Standard Desalting | GACGGGCGGTGWGTRCA | Dueholm et. al., 2022 ^6^ |
| ddPCR-515F-FAM | Standard Desalting | (FAM)-TGYCAGCMG-(ZEN)-CCGCGGTAA-(IBFQ) | Parada et al., 2016 ^7^ |
| **ssUMI workflow** | | | |
| ssUMI-8F-UMI | PAGE | GTATCGTGTAGAGACTGCGTAGG NNNYRNNNYRNNNYRNNNA GRGTTYGATYMTGGCTCAG | Callahan et. al., 2019 ^5^  Karst et. al., 2021 ^2^  ONT ^8^ |
| ssUMI-1391R-UMI | PAGE | AGTGATCGAGTCAGTGCGAGTG NNNYRNNNYRNNNYRNNN GACGGGCGGTGWGTRCA | Dueholm et. al., 2022 ^6^  Karst et. al., 2021 ^2^  ONT ^8^ |
| ssUMI-Universal-F | Standard Desalting | GGTGCTGAAGAAAGTTGTCGGTGTCTTTGTGTTAACCGTATCGTGTAGAGACTGCGTAGG | ONT ^8^ |
| ssUMI-Universal-R | Standard Desalting | GGTGCTGAAGAAAGTTGTCGGTGTCTTTGTGTTAACCAGTGATCGAGTCAGTGCGAGTG | ONT ^8^ |

**Table S11.** PCR thermocycling conditions for UMI-tagging (PCR1) and amplification (PCR2) for the ssUMI workflow.

| **Step** | **Temperature** | **Ramp Rate** | **Time** | **Cycles** |
| --- | --- | --- | --- | --- |
| **ssUMI-PCR1** | | | | |
| Initial denaturation | 98°C | max | 3 min | 1 |
| Denaturation  Annealing  Extension | 98°C  Touchdown from 66°C to 60°C  72°C | max  0.2°C/sec  max | 30 sec  90 sec  3 min | 2 |
| Final extension | 72°C | max | 5 min | 1 |
| Hold | 4°C | - | - | - |
| **ssUMI-EarlyPCR2** | | | | |
| Initial denaturation | 98°C | max | 3 min | 1 |
| Denaturation  Annealing  Extension | 98°C  Touchdown from 70°C to 63°C  72°C | max  0.2°C/sec  max | 20 sec  45 sec  3 min 30 sec | 5 |
| Denaturation  Extension | 98°C  72°C | max  max | 20 sec  4 min | 5 |
| Final extension | 72°C | max | 5 min | 1 |
| Hold | 4°C | - | - | - |
| **ssUMI-LatePCR2** | | | | |
| Initial denaturation | 98°C | max | 3 min | 1 |
| Denaturation  Extension | 98°C  72°C | max  max | 20 sec  4 min | 15 |
| Final extension | 72°C | max | 5 min | 1 |
| Hold | 4°C | - | - | - |

**Table S12.** Summary of ASVs for the ZymoBIOMICS Gut Microbiome Standard that did not perfectly match a sequence in the reference database, but matched the same species in the NCBI nr database using BLASTn. These were considered true-positives (i.e. error-free sequences). Note, this did not occur with 97% OTUs, only ASVs.

| **Feature** | **Matched Genome** | **Query Cover** | **E- Value** | **Percent Identity** | **Accession** |
| --- | --- | --- | --- | --- | --- |
| *ssUMI_std (ASVs)* | | | | | |
| Zotu47 | Escherichia coli ATCC 8739 chromosome, complete genome | 100.0% | 0 | 100.0% | CP022959.1 |
| Zotu53 | Escherichia coli strain NEB5-alpha_F'Iq chromosome, complete genome | 100.0% | 0 | 100.0% | CP053607.1 |
| Zotu55 | Escherichia sp. strain Esraa 4 16S ribosomal RNA gene, partial sequence | 100.0% | 0 | 100.0% | MT647245.1 |
| *ssUMI_rapid (ASVs)* | | | | | |
| Zotu47 | Escherichia coli ATCC 8739 chromosome, complete genome | 100.0% | 0 | 100.0% | CP022959.1 |
| Zotu54 | Escherichia coli strain NEB5-alpha_F'Iq chromosome, complete genome | 100.0% | 0 | 100.0% | CP053607.1 |
| Zotu56 | Escherichia sp. strain Esraa 4 16S ribosomal RNA gene, partial sequence | 100.0% | 0 | 100.0% | MT647245.1 |
| *Quality-filtered Raw Nanopore Reads (ASVs)* | | | | | |
| Zotu50 | Clostridioides difficile strain FDAARGOS_723 chromosome, complete genome | 100.0% | 0 | 100.0% | CP046327.1 |
| Zotu60 | Escherichia coli strain CFS3313 chromosome, complete genome | 100.0% | 0 | 100.0% | CP026939.2 |
| Zotu65 | Escherichia coli strain S17-1 chromosome, complete genome | 100.0% | 0 | 100.0% | CP040667.1 |
| Zotu67 | Escherichia sp. strain Esraa 4 16S ribosomal RNA gene, partial sequence | 100.0% | 0 | 100.0% | MT647245.1 |

**Table S13.** Summary of ASVs and OTUs for the two microbiome standards that did not perfectly match a sequence in the reference database, but matched (using BLASTn) to an organism in the NCBI nr database that was not present in the Microbiome Standard at over 97% identity. These were considered contaminants, and were filtered from downstream analyses.

| **Feature Type** | **Microbiome Standard** | **Data Mode** | **Feature** | **Matched Genome** | **Query Cover** | **E- Value** | **Percent Identity** | **Accession** |
| --- | --- | --- | --- | --- | --- | --- | --- | --- |
| ASV | DNA Standard | ssUMI_std | Zotu28 | Caballeronia mineralivorans strain PAMC 27316 | 100.0% | 0 | 100.00% | MT555326.1 |
| ASV | DNA Standard | ssUMI_rapid | Zotu28 | Caballeronia mineralivorans strain PAMC 27316 | 100.0% | 0 | 100.00% | MT555326.1 |
| OTU | DNA Standard | ssUMI_std | OTU9 | Caballeronia mineralivorans strain PAMC 27316 | 100.0% | 0 | 100.00% | MT555326.1 |
| OTU | DNA Standard | ssUMI_std | OTU10 | Oligoflexia bacterium strain STP885.7 | 100.0% | 0 | 99.93% | MT827150.1 |
| OTU | DNA Standard | ssUMI_rapid | OTU9 | Caballeronia mineralivorans strain PAMC 27316 | 100.0% | 0 | 100.00% | MT555326.1 |
| OTU | DNA Standard | ssUMI_std | OTU10 | Oligoflexia bacterium strain STP885.7 | 100.0% | 0 | 99.93% | MT827150.1 |
| OTU | Gut Standard | ssUMI_std | OTU13 | Caballeronia mineralivorans strain PAMC 27316 | 100.0% | 0 | 100.00% | MT555326.1 |
| OTU | Gut Standard | ssUMI_std | OTU15 | Oligoflexia bacterium strain STP885.7 | 100.0% | 0 | 99.93% | MT827150.1 |
| OTU | Gut Standard | ssUMI_std | OTU16 | Uncultured bacterium clone ncd411g11c1 | 100.0% | 0 | 97.58% | HM323381.1 |
| OTU | Gut Standard | ssUMI_rapid | OTU12 | Caballeronia mineralivorans strain PAMC 27316 | 100.0% | 0 | 100.00% | MT555326.1 |
| OTU | Gut Standard | ssUMI_rapid | OTU15 | Uncultured bacterium clone ncd411g11c1 | 100.0% | 0 | 97.58% | HM323381.1 |

**Table S14.** Full-length 16S rRNA gene ddPCR thermocycling conditions.

| **Step** | **Temperature** | **Ramp Rate** | **Time** | **Cycles** |
| --- | --- | --- | --- | --- |
| Enzyme Activation | 95°C | 2°C/sec | 10 min | 1 |
| Denaturation  Annealing  Extension | 94°C  60°C  72°C | 2°C/sec | 30 sec  1 min  4 min | 50 |
| Enzyme Deactivation | 98°C | 2°C/sec | 10 min | 1 |
| Hold | 4°C | - | - | - |

**Table S15.** Summary of UMI-tagging efficiencies of E. coli genomic DNA, estimated using ddPCR on full-length 16S rRNA gene copies of input material and on the product after 2-cycles of ssUMI-PCR1 (see Supporting Text).

| **Input *E. coli* 16S rRNA (copies)** | **UMI-tagging efficiency** |
| --- | --- |
| 1,000,000 | 30.5 ± 5.% |
| 500,000 | 44.3 ± 4% |
| 100,000 | 38.6 ± 2% |
| 50,000 | 37.6 ± 1% |
| 10,000 | 36.8 ± 3% |
